# *Rtel1* in hypothalamic oxytocin neurons regulates oxytocin output and social behaviors in mice

**DOI:** 10.64898/2026.09.10.750776

**Authors:** Junwen Wang, Xiaqing Wang, Zilong Qiu

## Abstract

RTEL1, a DNA helicase that resolves G-quadruplexes to maintain telomere and genome stability, has emerged as a candidate autism spectrum disorder risk gene, but its neural functions remain unknown. Here, we show that *Rtel1* haploinsufficiency in mice impairs social novelty and increases marble burying and anxiety-related behaviors without affecting locomotion or spatial learning. RTEL1 is expressed in oxytocin (OXT) neurons of the hypothalamic paraventricular nucleus, where *Rtel1* haploinsufficiency reduces OXT abundance and blunts social-evoked OXT sensor responses. RTEL1 associates with a G-quadruplex-forming sequence near the *Oxt* transcription start site, and purified RTEL1 binds and unwinds this structure in an ATP-dependent manner. *Rtel1* reduction in *Oxt*-lineage cells recapitulates the social and OXT phenotypes, whereas adult RTEL1 re-expression in paraventricular OXT neurons increases OXT abundance and ameliorates behavioral abnormalities. These findings uncover a non-telomeric function for RTEL1 linking promoter DNA structure to hypothalamic oxytocin output and social behavior.

## Introduction

Autism spectrum disorder (ASD) is a heterogeneous neurodevelopmental condition defined by defects in social communication and interaction together with restricted or repetitive behaviors and interests. Current U.S. estimates indicate ASD in approximately 2.3% of 8-year-old children[1,2] and 2.2% of adults[1]. Earlier detection and biologically informed stratification may improve access to individualized support and intervention[3], motivating efforts to identify risk genes and determine how they alter neural function. In a previous study of Chinese ASD probands, we identified candidate risk genes, including nine genes not represented in the SFARI Gene database at the time of analysis and recurrent variants in the helicase-encoding gene *RTEL1* among probands with IQ scores above 70 (FDR < 0.1) [4]. Variants in *RTEL1* have also been reported in individuals with other neurodevelopmental disorders[5,6]. Whether reduced RTEL1 dosage influences ASD-relevant neural circuits and behavior, however, remains unknown.

RTEL1 is an ATP-dependent DNA helicase essential for telomere maintenance and genome integrity[7]. Its modular architecture—including the helicase core and C-terminal interaction domains—is conserved across mammals[7,8]. RTEL1 dismantles displacement-loop intermediates, including telomeric loops, and resolves DNA secondary structures such as G-quadruplexes (G4s)[9,10]. These activities are required for efficient telomere replication: failure to unwind t-loops and telomeric G4s promotes telomere loss[11,12]. More broadly, RTEL1 deficiency causes replication defects and genome instability associated with unresolved G4 and R-loop structures[10,13,14].

Biallelic *RTEL1* variants cause Hoyeraal–Hreidarsson syndrome, a severe disorder characterized by bone-marrow failure, growth restriction, and neurological abnormalities including microcephaly, cerebellar hypoplasia, and intellectual disability[15–17]. In *C. elegans*, loss of the *rtel-1* ortholog causes genome-instability phenotypes and synthetic lethality with defects in other replication or repair pathways[18], whereas homozygous *Rtel1* disruption causes embryonic lethality in mice[19]. Although these phenotypes establish an essential role for RTEL1 in proliferating cells, RTEL1-dependent G4 processing can also influence transcription[20]. Indeed, *Rtel1* loss and pharmacological G4 stabilization elicit overlapping transcriptional changes in mouse embryonic fibroblasts[20], raising the possibility that RTEL1 has dosage-sensitive, non-telomeric functions in differentiated cells.

G4s are noncanonical four-stranded structures formed by guanine-rich sequences[21]. Four guanines assemble into planar G-quartets through Hoogsteen hydrogen bonds, and stacking of two or more quartets stabilizes the G4 fold[22]. Once thought to be largely restricted to telomeres, G4-forming sequences are abundant throughout mammalian genomes[23–25] and participate in DNA replication[26] and genome-stability control[27]. They are also enriched at mammalian gene promoters[24,28]; computational surveys predict at least one promoter G4 motif for more than 40% of human genes[29]. Promoter G4s can modulate transcription in a sequence- and context-dependent manner[30,31]. Studies of promoter G4s, their binding partners, and G4-interacting ligands illustrate multiple routes by which these structures can influence transcription[32–35]. These effects need not be uniformly repressive[30,31,34]. Consistent with a role for RTEL1 in this regulatory layer, *Rtel1*-null fibroblasts show telomere-independent transcriptional changes that preferentially affect genes bearing promoter G4-forming sequences and overlap with changes induced by the G4-interacting ligand TMPyP4[21]. These observations suggest that RTEL1 could couple promoter DNA structure to cell-type-specific gene expression, but this possibility has not been tested in neural circuits relevant to ASD.

Oxytocin (OXT) is a cyclic nonapeptide synthesized primarily by hypothalamic neurons in the paraventricular nucleus (PVN) and supraoptic nucleus (SON)[36,37]. OXT has peripheral endocrine actions[36,38] and is also delivered to distributed brain regions by central projections from PVN neurons[39,40], where it modulates social-cue processing, affect, and interaction[41,42]. Social encounters recruit PVN OXT neurons[43,44]. Across distinct circuits, OXT signaling contributes to medial amygdala-dependent social recognition[45], the formation of cortical representations of familiar individuals[46], PVN-to-ventral tegmental area social reward[47], and supraoptic nucleus-to-lateral septum regulation of social fear in lactating mice[48]. OXT further modulates olfactory bulb processing[49], anterior olfactory nucleus-dependent social recognition[50], and hippocampal social discrimination[51]. Its effects remain strongly context dependent: OXT signaling shapes territorial recognition, scent marking, and aggression[52–54], and projection-, cell-type-, and sex-specific studies reveal divergent effects on aggressive and defensive behavior[55–57]. Preclinical studies also implicate OXT in anxiety-related states[58,59], including responses to chronic isolation[60]. In adolescents, salivary OXT is correlated with separation-anxiety symptoms[61], and sleep-restriction studies implicate PVN OXT-neuron activity in anxiety-like behavior[62]. OXT has also been proposed as a target for canine separation anxiety, although efficacy remains uncertain[63]. Thus, OXT provides a plausible link between hypothalamic gene regulation and social and affective behavior, but the endogenous mechanisms governing *Oxt* expression remain incompletely defined.

Here, we asked whether *Rtel1* haploinsufficiency disrupts ASD-relevant behavior through a promoter-G4 mechanism in the hypothalamic OXT system. Global *Rtel1* heterozygous mice showed impaired social novelty and abnormalities in assays of repetitive and anxiety-like behavior. RTEL1 localized to OXT neurons in the PVN and associated with a G4-forming sequence near the *Oxt* transcription start site, and biochemical assays showed that RTEL1 unwinds this G4 substrate in vitro. *Rtel1* haploinsufficiency reduced OXT abundance and altered socially evoked OXT dynamics in the PVN. Conditional *Rtel1* haploinsufficiency in the *Oxt* lineage reproduced key behavioral abnormalities, whereas PVN-targeted *Rtel1* re-expression increased OXT levels and partially rescued behavior. Together, these findings link RTEL1 dosage to hypothalamic OXT biology and support a model in which impaired processing of an *Oxt* promoter G4 may contribute to altered social behavior.

## Results

### ASD-associated RTEL1 variants reduce protein abundance, and *Rtel1* haploinsufficiency alters OXT and PV expression

In our previous whole-exome sequencing study of Chinese individuals with autism spectrum disorder (ASD), we identified two *RTEL1* variants, the nonsense variant p.W289* and the missense variant p.R734W[4]. Mapping these variants onto the 996-amino-acid RTEL1 protein showed that p.W289* lies within the N-terminal helicase region, whereas p.R734W is located in the HNL1 domain near the G4-binding region (Figure 1A). Additional *RTEL1* variants have been reported in individuals with ASD, developmental delay, or other neurodevelopmental phenotypes[5,6]. To determine how the two ASD-associated variants affect protein abundance, we expressed FLAG-tagged wild-type and mutant RTEL1 constructs in HEK293T cells. Full-length FLAG-tagged p.W289* protein was undetectable, whereas the abundance of p.R734W RTEL1 was markedly reduced relative to wild-type RTEL1 (Figure 1B). Thus, both variants reduced the detectable abundance of full-length RTEL1 in this heterologous expression system.

**Figure 1.**
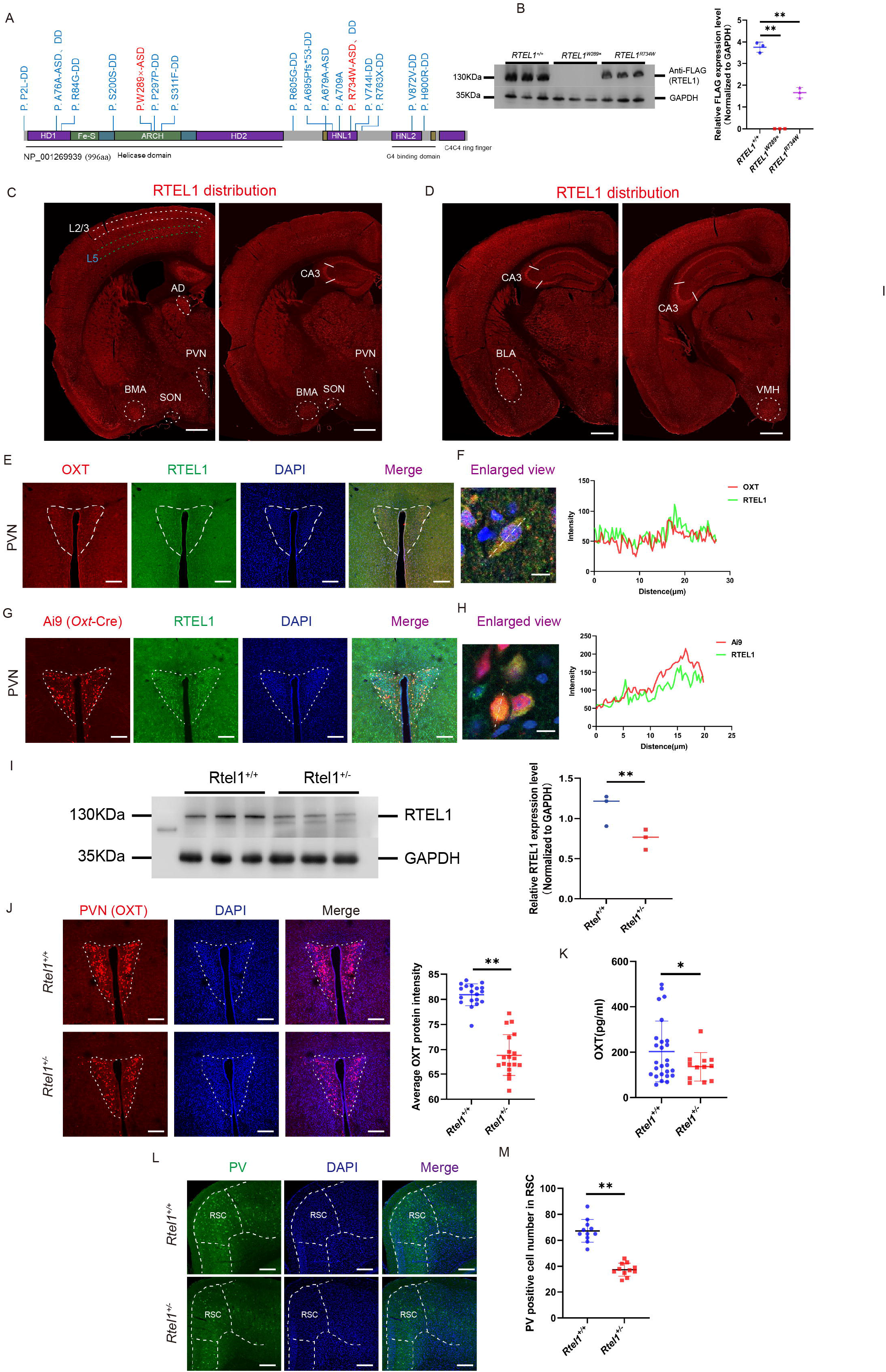
RTEL1 is broadly expressed in the adult mouse brain and in OXT neurons, and *Rtel1* haploinsufficiency reduces PVN OXT immunoreactivity. (A) Domain organization of human RTEL1 isoform NP_001269939 (996 amino acids). Previously reported variants associated with developmental disorders are shown in blue, and the de novo p.W289* and p.R734W variants identified in ASD probands are shown in red. HD, helicase domain; HNL, harmonin-N-like domain. (B) Representative immunoblot of FLAG-tagged wild-type RTEL1, RTEL1-p.W289*, and RTEL1-p.R734W expressed in HEK293T cells (left) and quantification of full-length FLAG-RTEL1 abundance normalized to GAPDH (right). No full-length FLAG-reactive product was detected for p.W289*, whereas p.R734W reduced the abundance of full-length RTEL1 relative to wild-type RTEL1 (n = 3 independent experiments). (C and D) Representative sagittal brain sections showing the distribution of RTEL1 immunoreactivity in the adult mouse brain at two mediolateral levels. RTEL1 signal was detected in cortical layers 2/3 and 5, hippocampal CA3, anterodorsal thalamic nucleus (AD), paraventricular hypothalamic nucleus (PVN), supraoptic nucleus (SON), basomedial amygdala (BMA), basolateral amygdala (BLA), and ventromedial hypothalamus (VMH). Scale bars, 500 μm. (E) Representative immunofluorescence images of the PVN stained for OXT (red), RTEL1 (green), and DAPI (blue), with merged channels. Scale bars, 200 μm. (F) Enlarged image of an OXT-positive cell expressing RTEL1 and representative line-scan fluorescence-intensity profiles showing the spatial overlap of OXT and RTEL1 signals. Scale bar, 10 μm. (G) Representative PVN sections from *Oxt*-Cre;Ai9 mice showing the Ai9 tdTomato *Oxt*-lineage reporter (red), RTEL1 immunoreactivity (green), DAPI (blue), and merged channels. Scale bars, 200 μm. (H) Enlarged image of tdTomato-positive Oxt-lineage cells expressing RTEL1 and representative line-scan fluorescence-intensity profiles for tdTomato and RTEL1. Scale bar, 10 μm. (I) Representative immunoblot of RTEL1 in whole-brain lysates from *Rtel1*^+/+^ and *Rtel1*^+/−^ mice (left) and quantification of RTEL1 abundance normalized to GAPDH (right; n = 3 mice per genotype). (J) Representative OXT immunofluorescence images of the PVN from *Rtel1*^+/+^ and *Rtel1*^+/−^ mice and quantification of mean PVN OXT fluorescence intensity. Nuclei were counterstained with DAPI. Multiple sections from n = 3 mice per genotype were analyzed. Scale bars, 200 μm (K) Serum OXT concentrations in *Rtel1*^+/+^ and *Rtel1*^+/−^ mice measured by ELISA (n = 9 mice per genotype). (L and M) Representative parvalbumin (PV) immunofluorescence images in the retrosplenial cortex (RSC) of *Rtel1*^+/+^ and *Rtel1*^+/−^ mice (L) and quantification of PV-immunoreactive cells within the outlined region (M). Nuclei were counterstained with DAPI. Scale bars, 200 μm Data are presented as mean ± SD. Statistical comparisons were performed using one-way ANOVA followed by Dunnett’s multiple-comparisons test in (B) and two-tailed unpaired Student’s t tests in (I), (J), (K), and (M). **p < 0.01; ns, not significant. See also Figure S1.

To examine the expression pattern of RTLE1 in the brain, we performed immunohistochemical experiments to the adult mouse brain. Immunostaining revealed broad RTEL1 expression throughout the brain, including the cerebral cortex, amygdala, and multiple hypothalamic neurons (Figures 1C, 1D and S1A).

Interestingly, RTEL1 immunoreactivity in CA3 neurons appeared predominantly cytoplasmic, whereas both nuclear and cytoplasmic signals were observed in cortical neurons (Figure S1B). Notably, RTEL1 was detected in nearly all OXT-immunoreactive and *Oxt*-lineage cells in the paraventricular nucleus (PVN), with substantial overlap also observed in the supraoptic nucleus (SON) (Figures 1E, 1F and S1C, S1D).

To validate RTEL1 expression in OXT neuronal populations independently of peptide immunostaining, we examined *Oxt*-Cre;Ai9 mice, in which cells with current or previous *Oxt*-Cre activity are permanently labeled with tdTomato. RTEL1 immunoreactivity extensively overlapped with Ai9-labeled cells in both the PVN and SON, as further supported by overlapping fluorescence-intensity profiles. In the PVN, nearly all OXT-immunoreactive cells and a comparable proportion of Ai9-labeled *Oxt* -lineage cells expressed RTEL1 (Figures 1G, 1H). In the SON, both approaches detected RTEL1 in approximately two-thirds of the respective cell populations, with no significant difference between the two labeling methods (Figure S1E, S1F). These results independently demonstrate broad RTEL1 expression in hypothalamic *Oxt* -lineage populations.

To examine the consequences of reduced *Rtel1* dosage *in vivo*, we used CRISPR–Cas9 to generate mice carrying a heterozygous deletion of approximately 32.5 kb within the *Rtel1* locus (Figure S1G). Western blotting of whole-brain lysates demonstrated significantly reduced RTEL1 protein abundance in *Rtel1*^+/−^ mice relative to wild-type littermates, confirming RTEL1 haploinsufficiency *in vivo* (Figure 1I). Adult *Rtel1*^+/−^ mice appeared grossly normal and showed no significant differences in body weight in either sex (Figures S1H, S1I). Their brains were grossly similar in appearance and showed no differences in weight (Figures S1J, S1K).

Consistent with an effect of reduced *Rtel1* dosage on the oxytocin system, *Rtel1*^+/−^ mice exhibited significantly reduced OXT immunoreactivity in the PVN (Figure 1J). Circulating OXT concentrations showed a lower mean in *Rtel1*^+/−^ mice, comparing to WT mice (Figure 1K). In addition, the number of PV-immunoreactive cells in the retrosplenial cortex was markedly reduced in *Rtel1*^+/−^ mice (Figures 1L and 1M). Together, these findings show that ASD-associated RTEL1 variants reduce the abundance of full-length RTEL1 and that *Rtel1* haploinsufficiency is accompanied by altered hypothalamic OXT and cortical PV expression.

### RTEL1 occupies a G4-forming region of the Oxt promoter and promotes ATP-dependent G4 resolution in vitro

Because *Rtel1* haploinsufficiency reduced OXT immunoreactivity in the PVN and RTEL1 can resolve G-quadruplex (G4) DNA, we investigated whether RTEL1 associates with a G4-forming regulatory sequence at the *Oxt* locus. In silico analysis of the 1-kb promoter regions of mammalian *Oxt* orthologs identified several putative G4-forming sequences (Figures 2A, 2B). Notably, positionally analogous G-rich motifs were located immediately upstream of the transcription start site (TSS) in humans (−32 to −12 bp), mice (−23 to −8 bp), and rats (−24 to −8 bp) (Figure 2A). We therefore focused on the proximal mouse sequence and designed primers amplifying a 135bp promoter fragment encompassing this candidate G4 motif (Figure 2B ).

**Figure 2.**
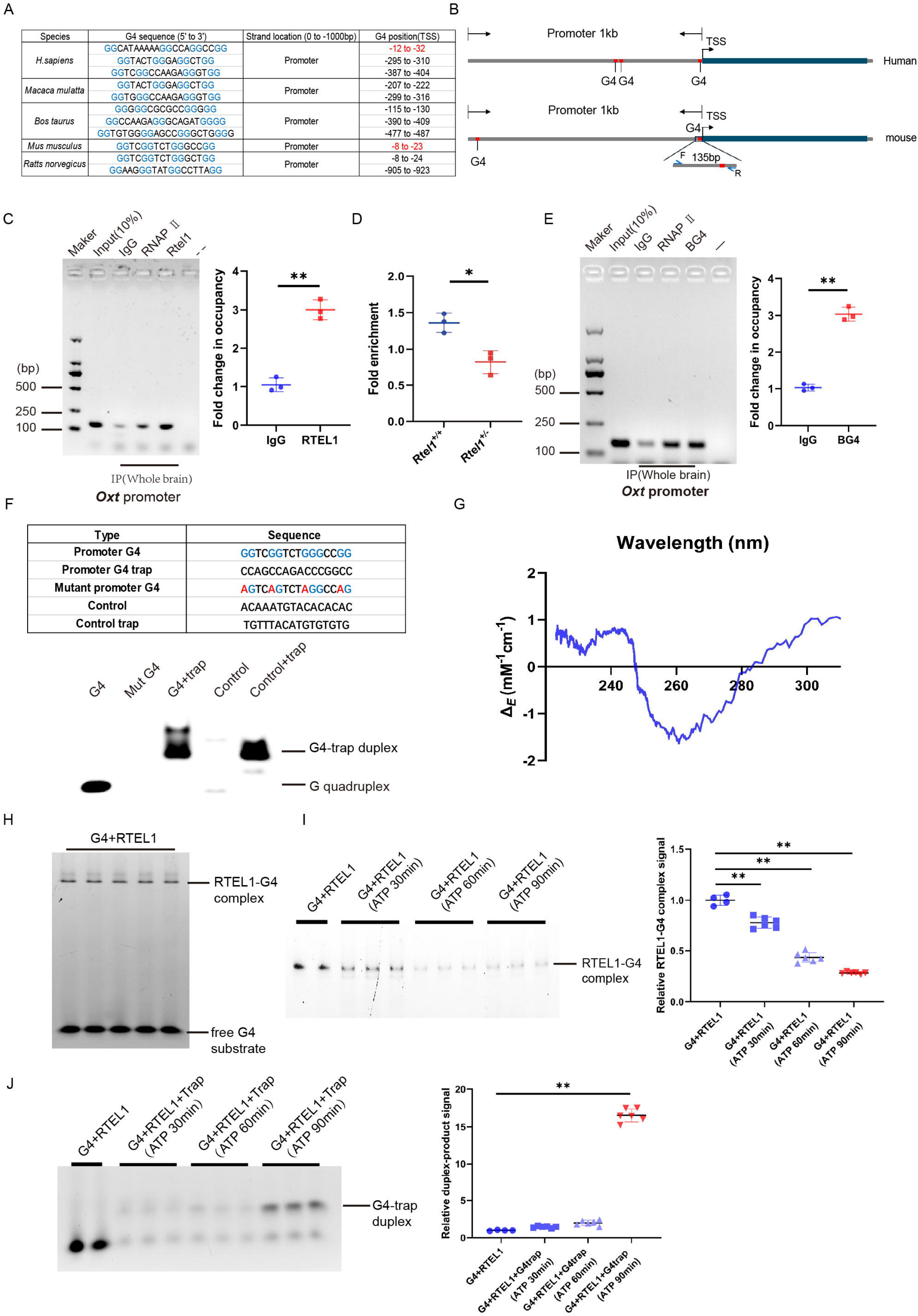
RTEL1 is enriched at a G4-containing region of the *Oxt* promoter and remodels the corresponding G4 substrate in vitro. (A) Predicted G-quadruplex-forming sequences within the 1-kb promoter regions of *OXT/Oxt* orthologs from the indicated mammalian species. Sequence positions are reported relative to the transcription start site (TSS). The TSS-proximal sequences in human and mouse are highlighted in red. (B) Schematic organization of predicted G4-forming sequences in the human *OXT* and mouse *Oxt* promoters. Forward (F) and reverse (R) primers amplify a 135-bp mouse promoter interval encompassing the TSS-proximal G4 motif. (C) Representative endpoint PCR analysis of the 135-bp *Oxt* promoter interval following chromatin immunoprecipitation (ChIP) from whole-brain chromatin using antibodies against RTEL1 or RNA polymerase II (RNA PII), or control IgG (left), and ChIP-qPCR quantification of RTEL1 enrichment relative to IgG (right; n = 3 independent ChIP experiments). Input represents 10% of the chromatin used for immunoprecipitation; “**−**” denotes the no-template PCR control. (D) RTEL1 ChIP-qPCR enrichment at the same proximal Oxt promoter interval in whole-brain chromatin from *Rtel1*^+/+^ and *Rtel1*^+/−^ mice (n = 3 independent samples per genotype). (E) Representative endpoint PCR analysis of the proximal *Oxt* promoter interval following ChIP from whole-brain chromatin using the G4-recognizing BG4 antibody, RNAPII antibody, or control IgG (left), and ChIP-qPCR quantification of BG4 enrichment relative to IgG (right; n = 3 independent ChIP experiments). Input represents 10% of the chromatin used for immunoprecipitation; “**−**” denotes the no-template PCR control. (F) Sequences of the wild-type *Oxt* promoter G4 oligonucleotide, its complementary trap strand, the G4-disrupting mutant, a length-matched control oligonucleotide, and the corresponding control trap strand (top). Native PAGE analysis of the indicated oligonucleotides following annealing under K^+^-containing conditions (bottom). The wild-type promoter oligonucleotide migrated as a G4 species, whereas annealing with its complementary trap strand generated a slower-migrating duplex product. (G) Circular dichroism spectrum of the *Oxt* promoter G4 oligonucleotide in buffer containing 150 mM K^+^. The positive signal near 295 nm and negative signal near 260 nm are consistent with a predominantly antiparallel G-quadruplex conformation. (H) Representative native gel-shift assay showing a slower-migrating species following incubation of the promoter G4 substrate with FLAG-tagged RTEL1, consistent with formation of an RTEL1–G4 complex. (I) Representative native gel (left) and densitometric quantification (right) of the RTEL1–G4 complex before and after incubation with ATP for 30, 60, or 90 min. The progressive reduction in complex signal is consistent with ATP-dependent remodeling or dissociation of the RTEL1–G4 complex. Values were normalized to the reaction without ATP. (J) Representative native gel (left) and quantification (right) of G4–trap duplex formation following the addition of ATP and the complementary trap strand to RTEL1–G4 reactions for 30, 60, or 90 min. The time-dependent accumulation of duplex product is consistent with G4 resolution followed by annealing of the trap strand. Data are presented as mean ± SD. Statistical significance was assessed using two-tailed Student’s t tests for the two-group comparisons in (C)–(E) and one-way ANOVA followed by Dunnett’s multiple-comparisons test in (I) and (J). *p < 0.05; **p < 0.01. See also Figure S2.

Chromatin immunoprecipitation (ChIP) from whole-brain tissue showed that an RTEL1 antibody recovered the proximal *Oxt* promoter fragment at significantly higher levels than the IgG control (Figure 2C). RTEL1 enrichment at this interval was reduced in *Rtel1^+/−^* mice relative to wild-type littermates, consistent with their reduced RTEL1 abundance (Figure 2D).

In an independent experiment, ChIP using BG4, an antibody that recognizes folded G4 structures, also enriched the same promoter fragment relative to IgG (Figure 2E). Thus, RTEL1 and a folded G4 structure were independently detected within the same promoter-proximal region of the endogenous *Oxt* locus.

We next examined whether the candidate sequence could form a G4 structure in vitro. Synthetic oligonucleotides were generated corresponding to the wild-type promoter sequence, a guanine-substituted mutant, a length-matched control, and their respective complementary trap strands (Figure 2F). Under potassium-containing conditions, the wild-type sequence formed a discrete species with the electrophoretic mobility expected for a folded G4. Annealing it with the complementary trap strand instead generated a slower-migrating duplex product (Figures 2F and S2A). The guanine-substituted sequence and length-matched control did not produce a comparable G4-associated species (Figure 2F). Circular dichroism spectroscopy further revealed a positive signal near 295 nm and a negative minimum near 260–270 nm, consistent with a predominantly antiparallel G4 conformation (Figure 2G).

To determine whether RTEL1 directly interacts with this structure, FLAG-tagged RTEL1 was expressed in HEK293T cells and isolated for biochemical analysis (Figure S2B). Electrophoretic mobility-shift assays revealed a slowly migrating RTEL1–G4 complex (Figure 2H). In contrast, the G4-disrupting mutant showed substantially reduced complex formation, indicating preferential RTEL1 binding to the intact G4-forming sequence (Figures S2C and S2D). Addition of ATP progressively reduced the G4-associated band over 30–90 min (Figure 2I). Moreover, in the presence of the complementary trap strand, ATP treatment resulted in time-dependent accumulation of duplex DNA, with the strongest signal observed after 90 min (Figure 2J). These results are consistent with ATP-dependent remodeling of the G4 by RTEL1, which exposes the G-rich strand and permits annealing to its complement.

Together, these chromatin and biochemical findings show that RTEL1 occupies a G4-containing region immediately upstream of the *Oxt* TSS and can preferentially bind and resolve the corresponding G4 substrate in vitro. These observations support a model in which RTEL1-dependent G4 resolution contributes to the maintenance of OXT abundance.

### Oxytocin sensor-based fiber photometry demonstrates diminished paraventricular oxytocin release in Rtel1 heterozygous knockout mice

Social engagement with conspecifics has been shown to activate oxytocin neurons in the paraventricular nucleus (PVN) and induce local oxytocin release. It has been confirmed by increases in activity to the social stimulus in PVN oxytocin neurons[43] and specifically upregulates Oxytocin release from PVN oxytocin neuron[55]. To further confirm the reduced OXT release in the PVN of Rtel1 heterozygous mice, we employed fiber photometry using the OT1.0 fluorescent sensor[64]. Firstly, we virally expressed the OT1.0 sensor in the PVN region of mice, and then observed changes in its fluorescence signal in social stimulation (Fig 3A, 3B). The experiments were conducted separately on six mice of different genotypes. During the experiment, the fluorescence signals within 10 seconds before and after social stimulation were recorded (Fig 3C, 3D). As shown in the experiment, after social stimulation, the OXT fluorescence signal in the PVN region of mice significantly increased overall (Fig 3E-G), although the sensitivity to social stimulation varied among individual mice. However, the OXT signal intensity in the PVN region of *Rtel1* gene heterozygous mice was significantly lower than that of the control group after social stimulation, indicating a lower release of OXT (Fig 3H).

**Figure 3.**
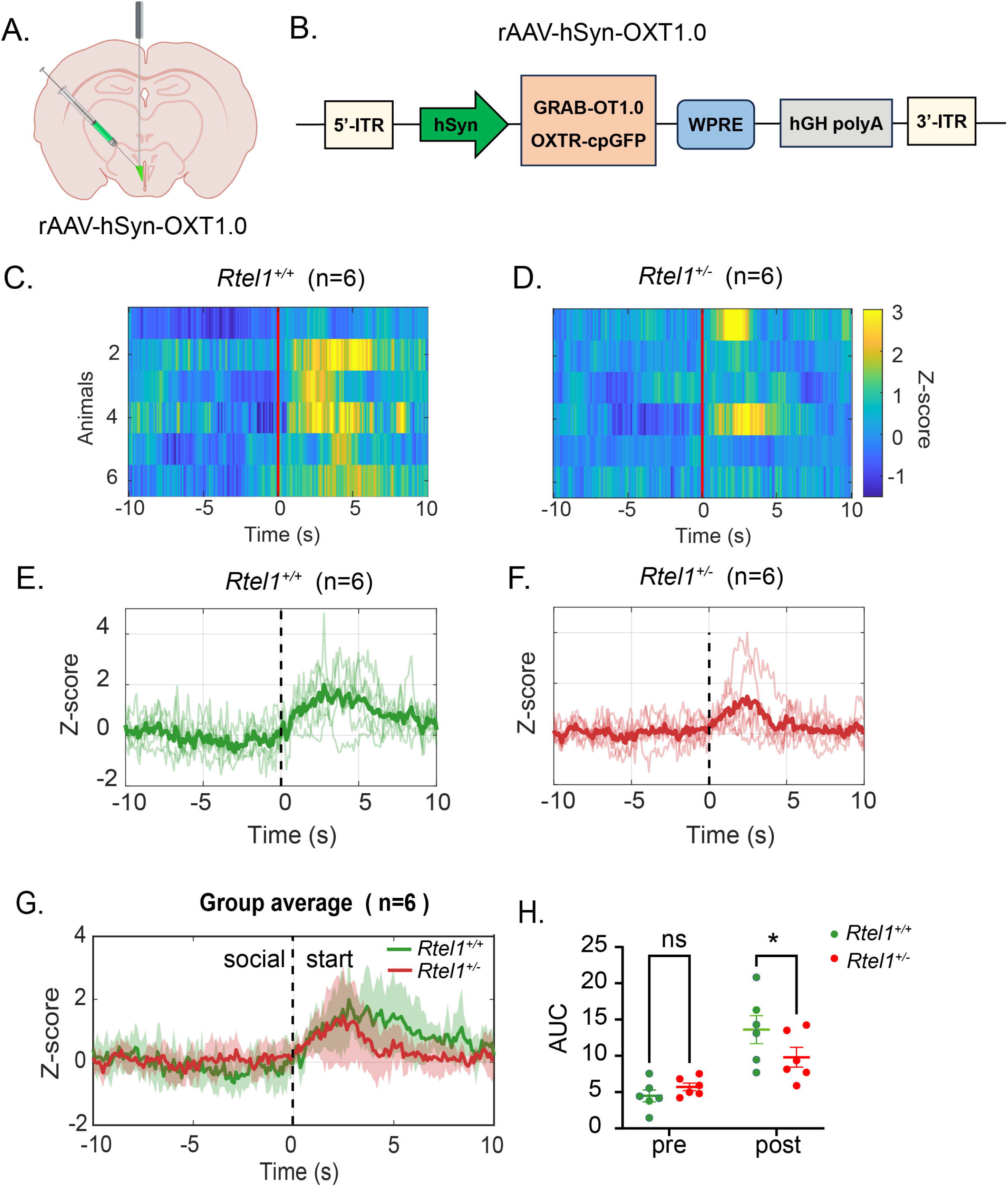
*Rtel1* haploinsufficiency attenuates social interaction–evoked OXT release in the PVN. (A) Schematic of viral delivery of the genetically encoded OXT sensor into the paraventricular nucleus (PVN) and optical-fiber implantation for fiber-photometry recording. (B) Schematic of the rAAV-hSyn-OXT1.0 expression cassette. The OXTR–circularly permuted GFP sensor (OXTR-cpGFP) was expressed under the human synapsin promoter. ITR, inverted terminal repeat; WPRE, woodchuck hepatitis virus post-transcriptional regulatory element; hGH poly(A), human growth hormone polyadenylation signal. (C and D) Heatmaps of z-score-normalized OXT1.0 fluorescence signals recorded from the PVN of individual *Rtel1*^+/+^ (C) and *Rtel1*^+/−^ (D) mice, aligned to the onset of social interaction at time 0. Each row represents one mouse (n = 6 mice per genotype). (E and F) Individual-animal traces (thin lines) and genotype-averaged traces (thick lines) of OXT1.0 fluorescence in *Rtel1*^+/+^ (E) and *Rtel1*^+/−^ (F) mice. Dashed vertical lines indicate the onset of social interaction. (G) Genotype-averaged OXT1.0 fluorescence responses aligned to social-interaction onset. Shaded regions indicate SEM (n = 6 mice per genotype). (H) Area under the curve (AUC) of the OXT1.0 fluorescence signal during the pre- and post-interaction epochs. Pre-interaction AUC did not differ between genotypes, whereas the social interaction–evoked response was attenuated in *Rtel1*^+/−^ mice. Each point represents one mouse. Data are presented as mean ± SEM. ns, not significant.

### *Rtel1* haploinsufficiency impairs social novelty and increases anxiety-related behavior

To determine the behavioral consequences of reduced *Rtel1* dosage, we subjected male *Rtel1*^+/−^ mice and their wild-type littermates to a battery of social, repetitive-like, anxiety-related, locomotor, and spatial-learning assays (Figure 4A) and examined the corresponding behaviors in an independent female cohort (Figure S3). In the sociability phase of the three-chamber test, male mice of both genotypes spent significantly more time investigating a conspecific than an empty enclosure, and their social-preference indices did not differ (Figures 4B and 4C). The same pattern was observed in females (Figures S3A and S3B), indicating preserved initial sociability in *Rtel1*^+/−^ mice. During the subsequent social-novelty phase, wild-type males preferentially investigated the novel conspecific over the familiar mouse, whereas *Rtel1*^+/−^ males showed no significant novelty preference and had a lower social-novelty index (Figures 4D and 4E). Female heterozygotes exhibited a comparable reduction in social-novelty preference (Figures S3C and S3D). Thus, *Rtel1* haploinsufficiency impaired preference for social novelty without disrupting initial social approach.

**Figure 4.**
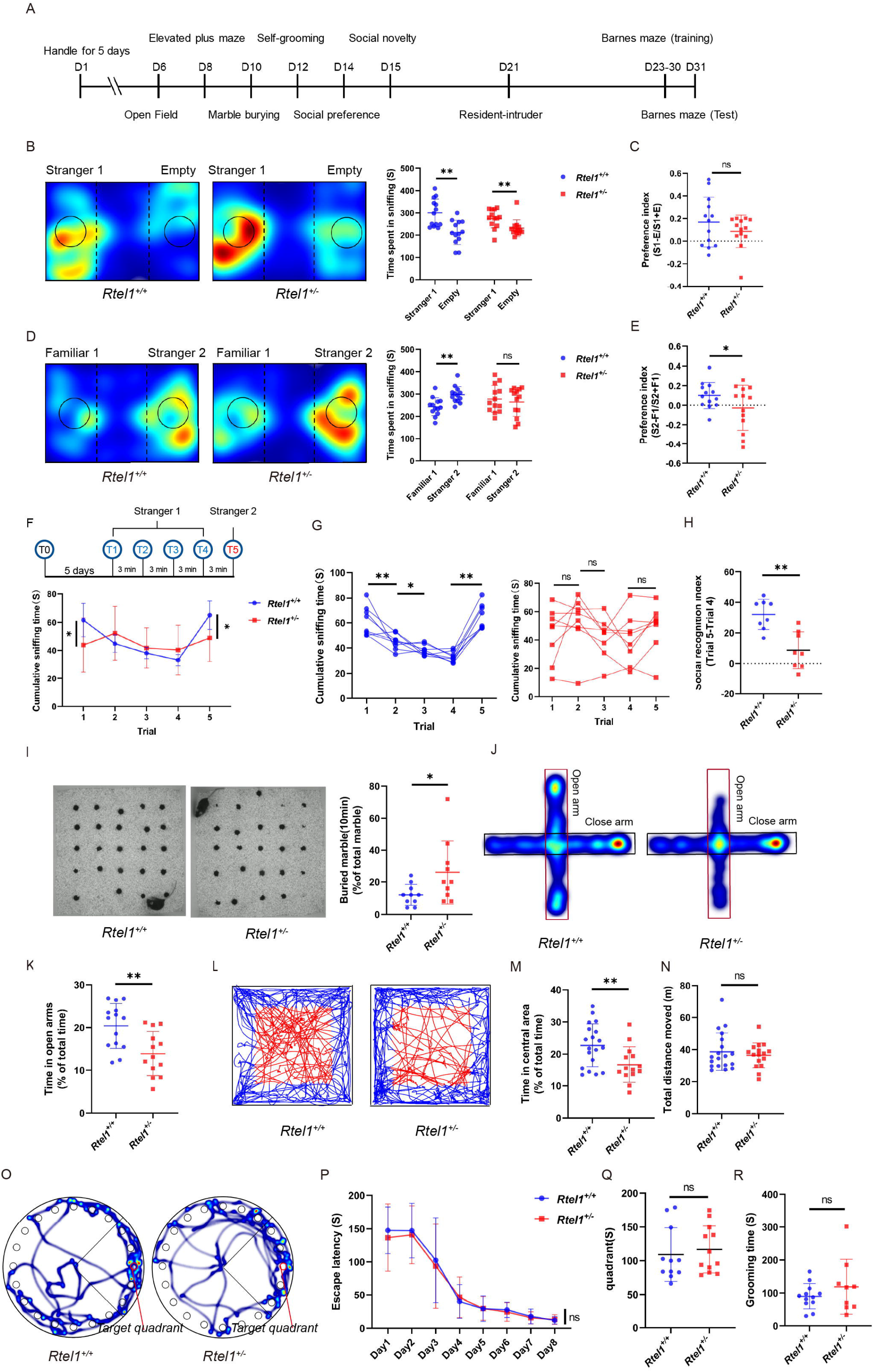
*Rtel1* haploinsufficiency impairs social novelty and increases marble burying and anxiety-related avoidance in male mice. (A) Experimental timeline for behavioral testing of male *Rtel1*^+/+^ and *Rtel1*^+/−^ mice. Mice underwent open-field, elevated-plus-maze, marble-burying, self-grooming, three-chamber social-preference and social-novelty, resident–intruder, and Barnes-maze tests on the indicated days. (B) Representative occupancy heatmaps during the sociability phase of the three-chamber test (left) and time spent sniffing stranger 1 (S1) or the empty enclosure (E) (right). Both genotypes spent more time investigating S1 than the empty enclosure. (C) Sociability preference index, calculated as (S1 − E)/(S1 + E). No significant genotype difference was detected. (D) Representative occupancy heatmaps during the social-novelty phase of the three-chamber test (left) and time spent sniffing the familiar mouse (F1) or a novel mouse (S2) (right). *Rtel1*^+/+^ mice preferentially investigated S2, whereas *Rtel1*^+/−^mice did not. (E) Social-novelty preference index, calculated as (S2 − F1)/(S2 + F1), showing a reduction in *Rtel1*^+/−^ mice. For (B)–(E), n = 13 mice per genotype. (F) Schematic of the resident–intruder habituation–dishabituation assay (top) and group-averaged cumulative sniffing time across five trials (bottom). Resident mice encountered the same intruder, stranger 1, during trials 1–4 and a novel intruder, stranger 2, during trial 5. n = 8 mice per genotype. (G) Individual cumulative sniffing times across the five resident–intruder trials, shown separately for *Rtel1*^+/+^ and *Rtel1*^+/−^ mice. Wild-type mice showed reduced investigation across repeated encounters with stranger 1 and renewed investigation of stranger 2, whereas these responses were attenuated in *Rtel1*^+/−^ mice. (H) Social-recognition index, calculated as the difference in sniffing time between trials 5 and 4, showing a reduction in *Rtel1*^+/−^ mice. For (F)–(H), n = 8 mice per genotype. (I) Representative images from the marble-burying test (left) and percentage of marbles buried during the 10-min test (right), showing increased marble burying in *Rtel1*^+/−^ mice. n = 10 mice per genotype. (J and K) Representative occupancy heatmaps from the elevated plus maze (J) and percentage of total test time spent in the open arms (K). *Rtel1*^+/−^ mice spent less time in the open arms than wild-type mice. n = 14 *Rtel1^+/+^* mice and 13 *Rtel1^+/−^* mice. (L–N) Representative open-field movement trajectories (L), percentage of total test time spent in the center area (M), and total distance traveled (N). *Rtel1*^+/−^ mice spent less time in the center, whereas total distance traveled was unaffected. n = 19 *Rtel1^+/+^*mice and 14 *Rtel1^+/−^* mice. (O–Q) Barnes-maze assessment of spatial learning and memory. Representative movement trajectories during the probe test, with the target quadrant indicated (O); escape latency across eight training days (P); and time spent in the target quadrant during the probe test following removal of the escape tunnel (Q). No significant genotype differences were detected during acquisition or the probe test. n = 11 *Rtel1^+/+^* mice and 12 *Rtel1^+/−^*mice. (R) Total self-grooming time. No significant genotype difference was detected. n = 12 *Rtel1^+/+^* mice and 9 *Rtel1^+/−^* mice. Data are presented as mean ± SD, and each symbol represents one mouse. Data in (B) and (D) were analyzed using two-way mixed-design ANOVA, with genotype as the between-subject factor and stimulus as the within-subject factor, followed by Šídák’s multiple-comparisons test. Resident–intruder data in (F) and (G) and Barnes-maze acquisition data in (P) were analyzed using two-way repeated-measures ANOVA followed by Šídák’s multiple-comparisons test where appropriate. Comparisons in (C), (E), (H), (I), (K), (M), (N), (Q), and (R) were performed using two-tailed unpaired Student’s t tests. *p < 0.05; **p < 0.01; ns, not significant. See also Figure S3.

The repeated resident–intruder assay provided convergent evidence of altered social recognition. Wild-type males progressively reduced their investigation of the same intruder over four successive trials and renewed their investigation when a novel intruder was introduced in trial 5. In contrast, *Rtel1*^+/−^ males showed neither clear habituation to the repeatedly presented intruder nor dishabituation to the novel intruder, resulting in a reduced trial 5–trial 4 social-recognition score (Figures 4F–4H). A similar pattern was observed in females (Figure S3E-3G). Heterozygous mice of both sexes also spent less time investigating the intruder during the first encounter, indicating that the resident–intruder phenotype was not restricted to the novelty trial. These findings suggest that reduced *Rtel1* dosage alters social habituation–dishabituation and social-novelty processing.

We next examined behaviors commonly used to assess repetitive-like phenotypes. Male *Rtel1*^+/−^ mice buried a greater proportion of marbles than wild-type males (Figure 4I). However, marble burying did not differ between female genotypes (Figures S3H and S3I), and self-grooming was unchanged in both male and female mice (Figure 4R; Figure S3J). Therefore, evidence for increased repetitive-like behavior was assay-specific and observed only in the male cohort.

In the elevated plus maze, male *Rtel1*^+/−^ mice spent less time in the open arms than wild-type littermates (Figures 4J and 4K). Female heterozygotes showed a similar reduction in elevated plus maze exploration (Figure S3K). In the open-field test, heterozygous mice of both sexes spent less time in the center, whereas total distance traveled was unchanged (Figures 4L–4N; Figures S3L and S3M). These results indicate increased anxiety-related avoidance without a detectable impairment in general locomotor activity.

Finally, both genotypes showed progressively shorter escape latencies during Barnes maze acquisition, with no detectable genotype effect on the learning curves of either male or female mice. During the probe trial, the time spent in the target quadrant was also comparable between genotypes (Figures 4O–4Q; Figure S3N). Thus, *Rtel1* haploinsufficiency did not measurably impair spatial learning or reference memory under these experimental conditions. Collectively, these results identify a behavioral profile characterized by impaired social-novelty processing and increased anxiety-related behavior in both sexes, together with increased marble burying observed only in males.

### *Rtel1* haploinsufficiency in *Oxt*-lineage cells impairs social novelty and increases anxiety-related behavior

To determine whether reduced RTEL1 dosage in oxytocin-expressing cells contributes to the behavioral alterations observed in germline *Rtel1*^+/−^ mice, we crossed *Oxt*-Cre mice with *Rtel1*^flox/+^ mice to generate *Oxt*-Cre; *Rtel1*^flox/+^ conditional heterozygotes. Cre-negative *Rtel1*^flox/+^ littermates served as controls. Because constitutive *Oxt* -Cre induces recombination in cells with current or previous *Oxt* expression, this manipulation targets *Oxt* -lineage cells rather than being anatomically restricted to the PVN. Male conditional heterozygotes underwent a battery of social, repetitive-like, anxiety-related, locomotor, and cognitive assays (Figure 5A), with corresponding behaviors examined in an independent female cohort (Figure S5).

**Figure 5.**
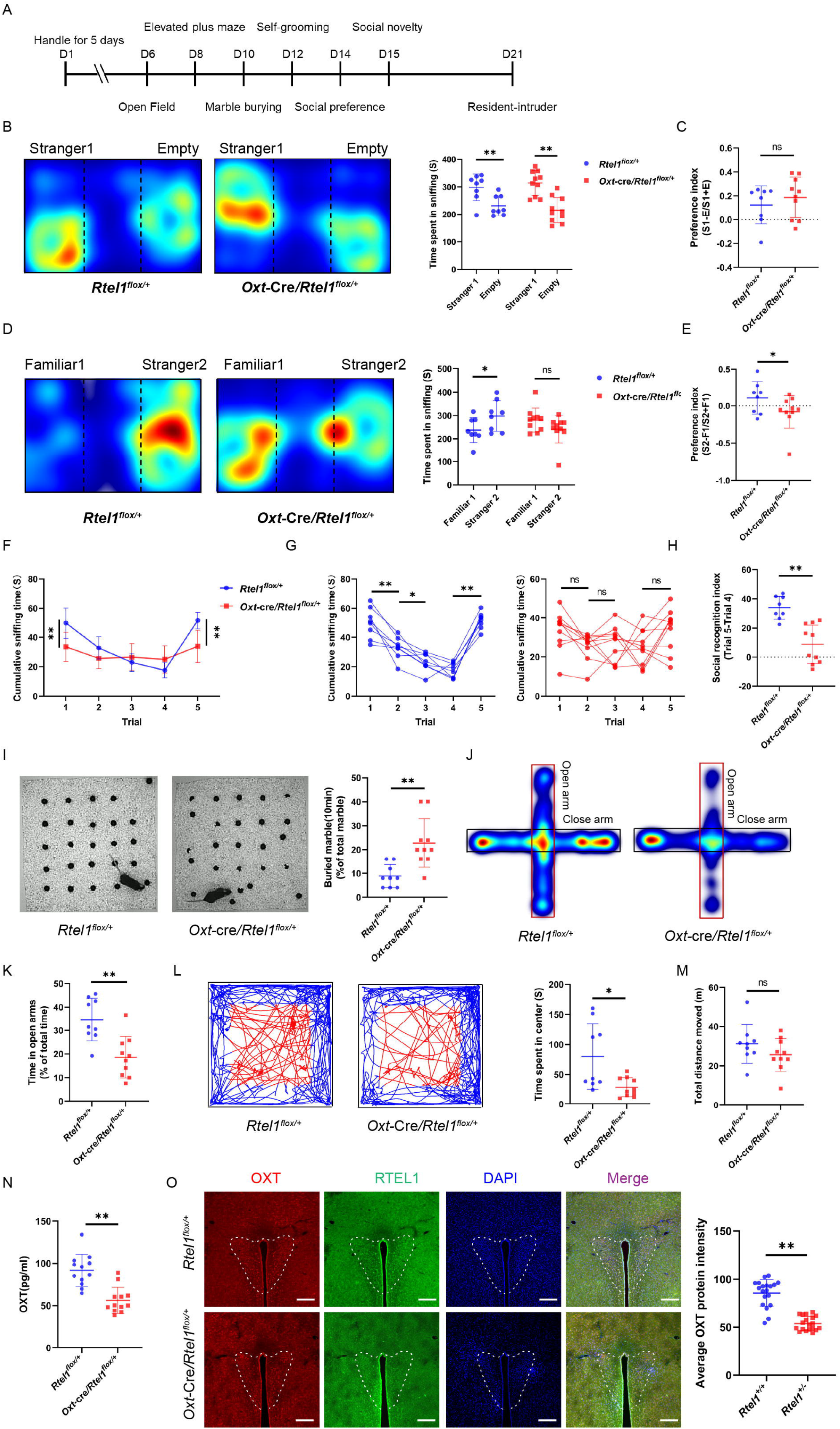
Heterozygous *Rtel1* deletion in *Oxt*-lineage cells impairs social novelty and reduces OXT abundance in male mice. (A) Experimental timeline for behavioral testing of male *Rtel1*^flox/+^ control and *Oxt*-Cre; *Rtel1*^flox/+^ mice. Mice underwent open-field, elevated-plus-maze, marble-burying, self-grooming, three-chamber social-preference and social-novelty, and resident–intruder tests on the indicated days. (B) Representative occupancy heatmaps during the sociability phase of the three-chamber test (left) and time spent sniffing stranger 1 (S1) or the empty enclosure (E) (right). Both genotypes spent more time investigating S1 than the empty enclosure. (C) Sociability preference index, calculated as (S1 − E)/(S1 + E). No significant genotype difference was detected. For (B)–(C), n = 8 *Rtel1*^flox/+^ control mice and 10 *Oxt*-Cre; *Rtel1*^flox/+^. (D) Representative occupancy heatmaps during the social-novelty phase of the three-chamber test (left) and time spent sniffing the familiar mouse (F1) or a novel mouse (S2) (right). Control mice preferentially investigated S2, whereas *Oxt*-Cre; *Rtel1*^flox/+^ mice did not show a significant preference. (E) Social-novelty preference index, calculated as (S2 − F1)/(S2 + F1), showing a reduction in *Oxt*-Cre; *Rtel1*^flox/+^ mice. For (D)–(E), n = 8 *Rtel1*^flox/+^ control mice and 10 *Oxt*-Cre; *Rtel1*^flox/+^. (F) Group-averaged cumulative sniffing time during the resident–intruder habituation–dishabituation assay. Resident mice encountered the same intruder, stranger 1, during trials 1–4 and a novel intruder, stranger 2, during trial 5. (G) Individual sniffing-time trajectories across the five resident–intruder trials, shown separately by genotype. Control mice showed reduced investigation across repeated encounters with stranger 1 and renewed investigation of stranger 2, whereas these responses were attenuated in *Oxt*-Cre; *Rtel1*^flox/+^ mice. (H) Social-recognition index, calculated as the difference in sniffing time between trials 5 and 4, showing a reduction in *Oxt*-Cre; *Rtel1*^flox/+^ mice. For (F)–(H), n = 8 *Rtel1*^flox/+^ control mice and 10 *Oxt*-Cre; *Rtel1*^flox/+^. (I) Representative images from the marble-burying test (left) and percentage of marbles buried during the 10-min test (right), showing increased marble burying in *Oxt*-Cre; *Rtel1*^flox/+^ mice. n = 8 *Rtel1*^flox/+^ control mice and 10 *Oxt*-Cre; *Rtel1*^flox/+^. (J and K) Representative occupancy heatmaps from the elevated plus maze (J) and percentage of total test time spent in the open arms (K). *Oxt*-Cre; *Rtel1*^flox/+^ mice spent less time in the open arms than controls. n = 8 *Rtel1*^flox/+^ control mice and 10 *Oxt*-Cre; Rtel1^flox/+^. (L and M) Representative open-field movement trajectories and time spent in the center area (L), and total distance traveled (M). *Oxt*-Cre; *Rtel1*^flox/+^ mice spent less time in the center, whereas total distance traveled was unchanged. For the behavioral analyses in (B)–(M), n = 9 *Rtel1*^flox/+^ control mice and 10 *Oxt*-Cre; *Rtel1*^flox/+^. (N) OXT concentrations measured by ELISA in serum samples obtained from *Rtel1*^flox/+^ and *Oxt*-Cre; *Rtel1*^flox/+^ mice. n = 7 mice each group. (O) Representative PVN sections stained for OXT (red), RTEL1 (green), and DAPI (blue), with merged channels, and quantification of mean PVN OXT fluorescence intensity. OXT immunoreactivity was reduced in *Oxt*-Cre; *Rtel1*^flox/+^ mice relative to controls. n = 3 mice per genotype. Scale bars, 200 μm. Data are presented as mean ± SD. Data in (B) and (D) were analyzed using two-way mixed-design ANOVA, with genotype as the between-subject factor and stimulus as the within-subject factor, followed by Šídák’s multiple-comparisons test. Resident–intruder data in (F) and (G) were analyzed using two-way repeated-measures ANOVA, with genotype as the between-subject factor and trial as the within-subject factor, followed by Šídák’s multiple-comparisons test. Between-genotype comparisons in (C), (E), (H), (I), and (K)–(N) were performed using two-tailed Welch’s t tests. Section-level measurements in (O) should be analyzed using a nested mixed-effects model with mouse included as the biological replicate. *p < 0.05; **p < 0.01; ns, not significant. See also Figures S4 and S5.

In the sociability phase of the three-chamber test, male mice of both genotypes spent significantly more time investigating a conspecific than an empty enclosure, and their social-preference indices did not differ (Figures 5B and 5C). In females, control mice preferred the conspecific, whereas conditional heterozygotes did not exhibit a significant within-group preference. However, the social-preference index did not differ significantly between female genotypes (Figure S5A), making the effect on initial female sociability equivocal. During the social-novelty phase, control males preferred the novel over the familiar conspecific, whereas conditional heterozygotes showed no significant novelty preference and had a lower novelty-preference index (Figures 5D and 5E). Female conditional heterozygotes similarly had a reduced novelty index and, unlike controls, spent more time investigating the familiar than the novel mouse (Figure S5B). Thus, heterozygous *Rtel1* deletion in *Oxt*-lineage cells consistently impaired social-novelty preference in both sexes.

The repeated resident–intruder assay provided convergent evidence of altered social investigation and recognition. Control males progressively reduced their investigation of the same intruder over four successive trials and renewed their investigation when a novel intruder was introduced in trial 5. Conditional heterozygous males showed neither clear habituation nor dishabituation and exhibited a lower trial 5–trial 4 social-recognition score (Figures 5F–5H). Female conditional heterozygotes showed a comparable loss of habituation–dishabituation and a reduced social-recognition score (Figure S5C and 5D). Conditional heterozygotes of both sexes also investigated the intruder less during the initial and novel-intruder trials, indicating that the resident–intruder phenotype was not restricted to the novelty comparison.

Male conditional heterozygotes buried more marbles than controls (Figure 5I), whereas self-grooming time was unchanged (Figure S4A). Neither marble burying nor self-grooming differed between female genotypes (Figures S5E and S5F). The evidence for increased repetitive-like behavior was therefore assay-specific and observed only in the male cohort. By contrast, anxiety-related avoidance was detected in both sexes. Male conditional heterozygotes spent less time in the open arms of the elevated plus maze and in the center of the open field, without a change in total distance traveled (Figures 5J–5M). Female conditional heterozygotes exhibited the same pattern (Figures S5G–S5I), indicating increased anxiety-related behavior without a detectable locomotor deficit.

During Barnes maze acquisition, escape latencies declined similarly across genotypes, and probe-trial occupancy of the target quadrant did not differ in either male or female mice (Figure S4B, S4C; Figures S5J, S5K). Thus, heterozygous Rtel1 deletion in *Oxt*-lineage cells did not measurably affect spatial learning or reference memory under these experimental conditions.

Finally, OXT concentrations measured by immunoassay were significantly reduced in male conditional heterozygotes (Figure 5N). Immunofluorescence analysis further demonstrated reduced OXT signal intensity in the PVN, accompanied by visibly diminished RTEL1 labeling (Figure 5O). Together, these results show that reducing *Rtel1* dosage in *Oxt*-lineage cells recapitulates the impaired social-novelty processing and increased anxiety-related behavior observed in germline *Rtel1*^+/−^ mice, together with increased marble burying in males. The accompanying reduction in OXT supports a functional relationship among RTEL1 dosage, oxytocin abundance, and these behavioral alterations but does not establish reduced OXT as their direct or sole cause.

### Adult RTEL1 re-expression in PVN Oxt-Cre^+^ neurons ameliorates behavioral abnormalities in *Rtel1* haploinsufficient mice

To determine whether restoring RTEL1 in adult PVN oxytocin neurons could ameliorate the behavioral consequences of *Rtel1* haploinsufficiency, we bilaterally injected two-month-old mice with AAV9-hSyn-DIO-mRtel1-3HA or the control AAV9-hSyn-DIO-GFP vector and began behavioral testing one month later. Three groups were compared: *Oxt*-Cre; *Rtel1*^+/+^ mice receiving DIO-GFP (WT-GFP), *Oxt*-Cre; *Rtel1*^+/−^ mice receiving DIO-GFP (mutant-GFP), and *Oxt*-Cre; *Rtel1*^+/−^ mice receiving DIO-mRtel1-3HA (mutant-RTEL1) (Figure 6A). This intersectional strategy was designed to restore RTEL1 selectively in *Oxt*-Cre^+^ neurons within the injected PVN.

**Figure 6.**
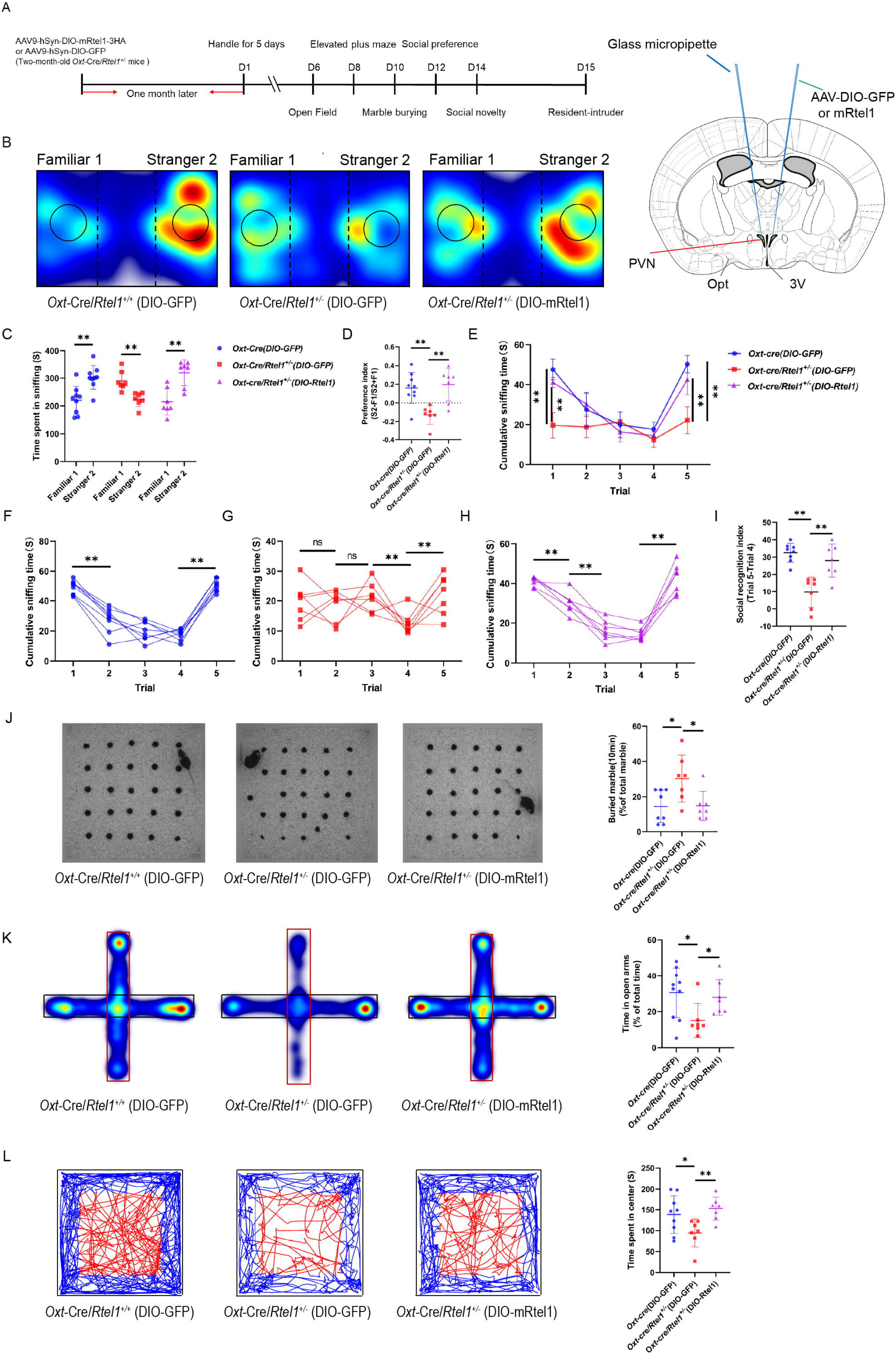
Adult PVN delivery of Cre-dependent RTEL1 ameliorates behavioral abnormalities in male *Rtel1* haploinsufficient mice. (A) Experimental design and behavioral-testing timeline (left). Two-month-old male mice received bilateral PVN injections of AAV9-hSyn-DIO-GFP or AAV9-hSyn-DIO-mRtel1-3HA. Behavioral testing began one month after injection (right). The three experimental groups were *Oxt*-Cre; *Rtel1*^+/+^ mice receiving DIO-GFP (WT-GFP), *Oxt*-Cre; *Rtel1*^flox/+^ mice receiving DIO-GFP (Het-GFP), and *Oxt*-Cre; *Rtel1*^flox/+^ mice receiving DIO-mRtel1-3HA (Het-RTEL1). (B) Representative occupancy heatmaps during the social-novelty phase of the three-chamber test for the three experimental groups. (C) Time spent sniffing the familiar mouse (F1) and a novel mouse (S2). WT-GFP and Het-RTEL1 mice preferentially investigated S2, whereas Het-GFP mice spent less time investigating S2 than F1. (D) Social-novelty preference index, calculated as (S2 − F1)/(S2 + F1). The index was reduced in Het-GFP mice relative to WT-GFP mice and increased in Het-RTEL1 mice relative to Het-GFP mice. For (B)-(D), n = 9 WT-GFP mice, n=7 Het-GFP and 7 Het-RTEL1. (E) Group-averaged cumulative sniffing time during the resident–intruder habituation–dishabituation assay. Resident mice encountered the same intruder, stranger 1, during trials 1–4 and a novel intruder, stranger 2, during trial 5. (F–H) Individual sniffing-time trajectories across the five resident–intruder trials for WT-GFP (F), Het-GFP (G), and Het-RTEL1 (H) mice. The habituation–dishabituation response was attenuated in Het-GFP mice and improved following DIO-mRtel1 delivery. (I) Social-recognition index, calculated as the difference in sniffing time between trials 5 and 4. The reduced index in Het-GFP mice was increased in Het-RTEL1 mice. For (E)-(I), n = 9 WT-GFP mice, n = 7 Het-GFP, and 7 Het-RTEL1 (J) Representative images from the marble-burying test (left) and percentage of marbles buried during the 10-min test (right). Increased marble burying in Het-GFP mice was reduced following DIO-mRtel1 delivery. n = 8 WT-GFP mice, n = 7 Het-GFP, and 7 Het-RTEL1 (K) Representative occupancy heatmaps from the elevated plus maze (left) and percentage of total test time spent in the open arms (right). Het-GFP mice spent less time in the open arms than WT-GFP mice, whereas DIO-mRtel1 delivery increased open-arm exploration. n = 9 WT-GFP mice, n = 7 Het-GFP, and 7 Het-RTEL1 (L) Representative open-field movement trajectories (left) and time spent in the center area (right). Center exploration was reduced in Het-GFP mice and increased following DIO-mRtel1 delivery. n = 9 WT-GFP mice, n = 7 Het-GFP, and 7 Het-RTEL1 Data are presented as mean ± SD, and each symbol represents one mouse. Data in (C) were analyzed using two-way mixed-design ANOVA, with treatment group as the between-subject factor and stimulus as the within-subject factor, followed by Šídák’s multiple-comparisons test. Resident–intruder data in (E) were analyzed using two-way repeated-measures ANOVA, with treatment group as the between-subject factor and trial as the within-subject factor, followed by Šídák’s multiple-comparisons test. Within-group trial comparisons in (F)–(H) were analyzed using repeated-measures ANOVA with multiplicity-adjusted comparisons. Data in (D) and (I)–(L) were analyzed using one-way ANOVA followed by Tukey’s multiple-comparisons test. *p < 0.05; **p < 0.01; ns, not significant. See also Figure S6.

During the social-novelty phase of the three-chamber test, WT-GFP males preferentially investigated the novel conspecific, whereas mutant-GFP males spent more time with the familiar mouse and exhibited a reduced novelty-preference index. Cre-dependent RTEL1 re-expression restored preferential investigation of the novel mouse and significantly increased the novelty-preference index relative to mutant-GFP animals (Figures 6B–6D). Female mice showed a comparable pattern: WT-GFP females preferred the novel conspecific, this preference was reversed in mutant-GFP females, and normal novelty preference was recovered following RTEL1 re-expression (Figures S6A–S6C). Thus, adult RTEL1 re-expression in PVN Oxt-Cre^+^neurons ameliorated impaired social-novelty preference in both sexes.

Consistent results were obtained in the repeated resident–intruder assay. WT-GFP males progressively reduced their investigation of the repeatedly presented intruder and renewed their investigation when a novel intruder was introduced in trial 5. Mutant-GFP males showed reduced investigation during the initial and novel-intruder trials and a lower trial 5–trial 4 social-recognition score. RTEL1 re-expression restored the habituation–dishabituation pattern and increased the social-recognition score relative to mutant-GFP males (Figures 6E–6I). Female mutant-GFP mice also showed an attenuated novelty-evoked response, whereas RTEL1 re-expression restored the increase in investigation during trial 5 and improved the social-recognition score (Figures S6D–S6H).

In the marble-burying assay, mutant-GFP males buried more marbles than WT-GFP controls, and RTEL1 re-expression significantly reduced this increase (Figure 6J). Marble burying and self-grooming were not examined in the female rescue dataset; therefore, rescue of repetitive-like behavior can be concluded only for marble burying in males.

RTEL1 re-expression also ameliorated anxiety-related avoidance. Mutant-GFP males spent less time in the open arms of the elevated plus maze and in the center of the open field than WT-GFP controls. Both measures were significantly increased by RTEL1 re-expression toward WT-GFP levels (Figures 6K and 6L). Equivalent improvements in open-arm and open-field-center exploration were observed in female mutant-RTEL1 mice (Figures S6I and S6J).

Collectively, these results demonstrate that adult RTEL1 re-expression in PVN Oxt-Cre^+^ neurons ameliorates social-novelty and social-recognition abnormalities and anxiety-related avoidance in male and female *Rtel1* haploinsufficient mice, while also reducing increased marble burying in males. These findings identify PVN *Oxt*-Cre^+^ neurons as a population in which restoring RTEL1 dosage can improve the behavioral consequences of Rtel1 haploinsufficiency. However, these experiments do not establish whether the behavioral improvements are mediated by restoration of OXT production.

## Discussion

Our findings identify a dosage-sensitive role for RTEL1 in hypothalamic oxytocin biology and social behavior. Two *RTEL1* variants identified in ASD probands without reported developmental delay or intellectual disability reduced the abundance of full-length RTEL1 when expressed in heterologous cells[4]. In mice, *Rtel1* haploinsufficiency impaired social-novelty preference and altered social habituation–dishabituation in both sexes, increased anxiety-related avoidance, and increased marble burying in males, while largely preserving initial sociability, locomotor activity, self-grooming, and spatial learning and memory. RTEL1 was broadly expressed in the adult mouse brain and was detected in most oxytocin (OXT) neurons of the paraventricular nucleus (PVN). Molecular analyses further linked RTEL1 to a G-quadruplex-forming sequence near the *Oxt* transcription start site. Heterozygous *Rtel1* deletion in Oxt-lineage cells reproduced the principal OXT and behavioral abnormalities, whereas adult Cre-dependent RTEL1 re-expression in PVN Oxt-Cre-positive neurons ameliorated the social and anxiety-related phenotypes. Together, these findings support a model in which RTEL1 dosage influences hypothalamic OXT function and selected social behaviors.

The human genetic evidence should nevertheless be interpreted cautiously. The p.W289* nonsense and p.R734W missense variants were identified in ASD probands in our previous cohort[4], and additional rare *RTEL1* variants have been reported in individuals with neurodevelopmental disorders[5,6]. In the present study, p.W289* produced no detectable full-length tagged RTEL1, whereas p.R734W reduced protein abundance. Because these experiments used ectopic cDNA expression, however, they do not reproduce endogenous transcription, nonsense-mediated decay, or cell-type-specific regulation. Moreover, the mouse model carries an approximately 32.5-kb heterozygous deletion within the *Rtel1* locus rather than the human p.W289* variant. It therefore models reduced *Rtel1* dosage but does not establish variant-specific pathogenicity. The combined human and mouse results strengthen the biological plausibility of *RTEL1* haploinsufficiency as a neurodevelopmental risk mechanism, but larger genetic cohorts, segregation or burden analyses, patient-derived cellular models, and functional evaluation of additional alleles will be required to establish the strength and phenotypic range of this association. Accordingly, *RTEL1* is more appropriately described as a candidate ASD-risk gene identified in probands without reported developmental delay or intellectual disability than as an established gene for “high-functioning autism.”

The behavioral profile of *Rtel1* haploinsufficiency was more selective than a generalized autism-like syndrome. Germline *Rtel1*^+/−^ mice of both sexes retained initial social approach but failed to exhibit the normal preference for a novel conspecific. Altered habituation and dishabituation during repeated resident–intruder encounters provided convergent evidence for disrupted social-recognition processing. However, reduced investigation during the first encounter suggests that changes in social motivation, olfactory processing, arousal, or anxiety could also contribute. Increased anxiety-related avoidance was consistently observed in the elevated plus maze and open field without a corresponding reduction in locomotion. By contrast, the repetitive-like phenotype was sex and assay dependent: marble burying was increased in males but not females, and self-grooming was unchanged in both sexes. Because marble burying can reflect digging or anxiety as well as repetitive behavior, this result should not be interpreted in isolation as evidence of stereotypy. Normal Barnes-maze acquisition and probe performance further argue against a broad spatial learning or memory deficit. Thus, the most reproducible behavioral consequence of *Rtel1* haploinsufficiency was impaired social-novelty processing accompanied by increased anxiety-related avoidance.

These findings also extend the known biology of RTEL1 beyond its canonical functions at replication forks and telomeres. RTEL1 is a conserved ATP-dependent helicase that supports genome-wide replication, counteracts telomeric G-quadruplexes, and promotes T-loop disassembly, thereby protecting telomeres from replication-associated instability[7,11,13,16,65]. Previous studies have also reported broad transcriptional alterations and increased G4- and R-loop-associated replication–transcription conflicts following RTEL1 loss[20]. Our results identify the *Oxt* promoter as a candidate neural substrate of a non-telomeric RTEL1 function. Positionally similar G-rich motifs were identified immediately upstream of the human, mouse, and rat *Oxt* transcription start sites. Chromatin immunoprecipitation with RTEL1 or the G4-recognizing BG4 antibody independently enriched the same promoter-proximal interval. The corresponding synthetic oligonucleotide adopted a G4-compatible conformation under potassium-containing conditions, while RTEL1 associated preferentially with the wild-type G4 substrate and promoted ATP-dependent substrate remodeling and duplex formation in vitro.

The present data support, but do not yet prove, direct RTEL1-dependent regulation of endogenous *Oxt* transcription through this promoter G4. RTEL1 and BG4 enrichment was measured in separate chromatin immunoprecipitation experiments using whole-brain tissue; these assays therefore do not demonstrate simultaneous RTEL1–G4 occupancy or establish that the interaction occurs specifically in PVN OXT neurons. Likewise, remodeling of a synthetic substrate in vitro does not demonstrate resolution of the endogenous promoter structure in intact neurons. Definitive tests would include selectively disrupting the endogenous G4-forming sequence, measuring nascent *Oxt* transcription and RNA polymerase occupancy, and comparing rescue by wild-type and helicase-deficient RTEL1. Cell-type-resolved chromatin analysis would further determine whether RTEL1 occupancy and G4 folding are altered specifically in OXT neurons. Such experiments are important because changes in OXT immunoreactivity can reflect altered synthesis, processing, storage, or release rather than transcription alone.

Several independent findings nevertheless connect RTEL1 dosage to the OXT system. RTEL1 was detected in most PVN OXT neurons, and germline *Rtel1* haploinsufficiency reduced PVN OXT immunoreactivity and blunted social-evoked OXT-sensor responses. Heterozygous deletion of *Rtel1* in *Oxt*-lineage cells similarly decreased OXT abundance and reproduced the social-novelty and anxiety-related phenotypes of germline heterozygotes. These results support an important contribution of RTEL1 within the OXT system. However, constitutive *Oxt*-Cre labels cells with current or previous *Oxt* expression in the PVN, supraoptic nucleus, and potentially peripheral tissues; this manipulation is therefore *Oxt*-lineage-specific rather than PVN-specific.

Bilateral delivery of a Cre-dependent RTEL1 vector to the adult PVN provided a more spatially restricted test of this model. RTEL1 re-expression ameliorated social-novelty preference, resident–intruder social-recognition measures, and anxiety-related avoidance in both sexes and reduced elevated marble burying in males. The efficacy of an adult intervention suggests that at least part of the phenotype remains modifiable after development and is not solely attributable to irreversible developmental abnormalities. Nevertheless, these experiments compared independently treated groups without pretreatment behavioral measurements and therefore demonstrate amelioration rather than reversal. Moreover, the rescue figures do not directly establish restoration of OXT abundance or demonstrate that OXT is the necessary downstream mediator. Direct restoration of OXT signaling, together with OXT-receptor blockade during RTEL1 rescue, would provide stronger causal and epistatic tests of the proposed RTEL1–OXT–behavior pathway. Quantitative confirmation of viral targeting and RTEL1 expression in the rescue cohorts would also strengthen the anatomical interpretation.

The neural findings should be considered within the broader context of RTEL1-associated telomere biology. Complete *Rtel1* loss causes early embryonic lethality in mice, whereas heterozygous animals are viable and grossly normal, consistent with previous genetic studies[19,66]. RTEL1 deficiency in humans is best established as a cause of telomere biology disorders, with severe neurodevelopmental manifestations particularly associated with biallelic pathogenic variants[15,67]. Monoallelic variants show variable expressivity and incomplete penetrance and should not be assumed to lack neurological consequences. The viable human cases generally carry alleles retaining some RTEL1 function and therefore cannot be directly compared with complete-null mouse embryos. Furthermore, the present deletion model was not evaluated for telomere length, replication stress, DNA damage, bone-marrow failure, or other telomere-biology phenotypes. Canonical genome-maintenance defects could consequently contribute to the neural phenotype alongside the proposed promoter-G4 mechanism.

Our observation of widespread RTEL1 immunoreactivity in the adult mouse brain also differs from an earlier bulk tissue survey in which brain expression was not detected.[19] Differences in age, assay sensitivity, tissue preparation, or cellular resolution could account for this discrepancy, and orthogonal validation at the transcript and single-cell levels would strengthen the conclusion. The reduction in parvalbumin-immunoreactive cells in the retrosplenial cortex further suggests that germline *Rtel1* haploinsufficiency may affect neural populations beyond the OXT system. Future studies using inducible, region- and cell-type-specific manipulations will be important for distinguishing developmental effects from ongoing functions of RTEL1 in the adult brain.

In summary, this study connects RTEL1 dosage to a promoter-proximal G-quadruplex, hypothalamic OXT biology, and social-novelty behavior. The convergence of germline haploinsufficiency, *Oxt*-lineage deletion, biochemical G4 assays, OXT measurements, and adult PVN-targeted complementation supports a non-telomeric neural function for RTEL1. Independent human genetic validation, endogenous interrogation of the Oxt promoter G4, and direct OXT epistasis experiments remain necessary to establish the proposed causal chain. RTEL1 should therefore be regarded as a candidate ASD and neurodevelopmental risk gene whose neural functions merit further investigation.

## Ethical considerations

All procedures were reviewed and approved by the Institutional Animal Care and Use Committee of Songjiang Hospital Affiliated to Shanghai Jiao Tong University School of Medicine under protocol “ACE-004-2025”.

## Acknowledgments

This work was supported by the following grants: National Science and Technology Major Project (2025ZD0214700), National Natural Science Foundation of China (#82430046 ZQ), and Project of Medical Technology Research and Transformation supported by Shanghai Municipal Health Commission (2024ZZ1007).

ChatGPT (OpenAI) was used to assist with language editing, manuscript organization, and drafting. The authors independently reviewed and verified all scientific content, analyses, interpretations, and references and accept full responsibility for the manuscript.

