## Supplementary material for "*Rtel1* in hypothalamic oxytocin neurons regulates oxytocin output and social behaviors in mice": Supp figures

**Materials and Methods**

**Supplemental Figures 1-6**

**Materials and Methods**

**Animals**

All mice were bred and maintained in accordance with the institutional animal care regulations of the Shanghai Laboratory Animal Center. Animals were housed in a specific pathogen‑free (SPF) facility on the 8th floor of the Songjiang Research Institute, Shanghai Jiao Tong University School of Medicine. Mice were provided with sterilized feed and drinking water, and were kept under controlled environmental conditions: room temperature at 20–25 °C, relative humidity of 30–70 %, and a 12/12‑h light/dark cycle (lights on from 07:00 to 19:00).

Rtel1 haploinsufficient mice were purchased from BIOCYTOGEN (https://biocytogen.com.cn/tool-mice) and generated by heterozygous knockout of the Rtel1 gene using CRISPR/Cas9 technology. Genotyping was performed with the following three primers: common forward primer, 5′‑CTGACCCTACAAGGACAGCTTCC‑3′; wild‑type reverse primer, 5′‑CCCAGTCTGGAATGGATGAGGGTAG‑3′; and mutant reverse primer, 5′‑GAGAGGCCACTAGGGCTTCAAAGTC‑3′.

Oxt‑Cre mice were obtained from The Jackson Laboratory and genotyped using the following primers: common forward primer, 5′‑TTTGCAGCTCAGAACACTGAC‑3′; wild‑type reverse primer, 5′‑AGCCTGCTGGACTGTTTTTG‑3′;

and mutant reverse primer, 5′‑ACACCGGCCTTATTCCAAG‑3′. All mouse strains were on a C57BL/6J genetic background.

**Behavioral Tests**

All behavioral tests were performed on mice aged 8 weeks. Experiments were conducted during the light phase to align with the animals’ circadian rhythm. Before testing, healthy littermates were selected whenever possible and subjected to a five‑day handling protocol to minimize fear and unfamiliarity toward the experimenter. Before each behavioral session, mice were transferred to the testing room and allowed to habituate for approximately 1 h to reduce anxiety, and the apparatus was thoroughly cleaned between subjects. The behavioral battery assessed anxiety‑like behavior, stereotyped behaviors, social interaction, and learning/memory, using the following paradigms: open field test, elevated plus maze, marble burying test, grooming test, three‑chamber social interaction test, resident–intruder test, and Barnes maze test.

For rescue experiments in *Oxt*‑Cre/*Rtel1^+/-^* mice, animals at 8 weeks of age received intracranial injections of AAV9‑hSyn‑DIO‑mRtel1‑3HA or AAV9‑hSyn‑DIO‑GFP (control)(PackGene (https://www.packgene.cn/)). Behavioral testing commenced four weeks after viral expression to allow sufficient transduction. The rescue battery focused on autism‑like phenotypes observed in Rtel1 haploinsufficient mice and included the open field test, elevated plus maze, marble burying test, three‑chamber social interaction test, and resident–intruder test. All behavioral data were analyzed using Noldus EthoVision XT software (<https://noldus.com/ethovision>).

**Open Field Test**

The open field test was used to assess anxiety‑like behavior and locomotor activity. The apparatus consisted of a white plastic box (0.75 cm thickness) with dimensions of 40 cm (length) × 40 cm (width) × 40 cm (height). Before each session, the arena was thoroughly cleaned to eliminate any residual odors, and the illumination level was set to 50 lux. At the start of the test, each mouse was placed in the center of the arena and allowed to explore freely for 10 min, during which its trajectory was recorded. After the session, the time spent in the central zone (24 cm × 24 cm) and the total distance traveled over the 10‑min period were analyzed.

**Elevated Plus Maze**

The elevated plus maze was used to assess anxiety‑like behavior in mice. The apparatus consisted of two open arms (30 × 5 × 1 cm) and two closed arms (30 × 5 × 15 cm), with the entire maze elevated 50 cm above the floor. Before testing, the apparatus was cleaned and the illumination was set to 50 lux. Each mouse was placed in the central area of the maze and allowed to explore freely for 6 min, during which its trajectory was recorded. The time spent in the open arms was subsequently analyzed. Data acquisition and analysis were performed using Noldus EthoVision XT software (https://noldus.com/ethovision), and graphical representation was carried out with GraphPad Prism 9.

**Marble Burying Test**

The marble burying test was performed in a white plastic box (40 × 40 × 40 cm). Prior to each trial, the apparatus was cleaned to remove any residual odors. Clean bedding was then placed into the box to a depth of 6 cm, and 25 black glass marbles were arranged on the surface in a 5 × 5 grid. Each mouse was placed in the box and allowed to explore freely for 10 min. After the session, the number of buried marbles was recorded; a marble was considered buried if more than 50 % of its surface was covered by bedding. Between trials, fecal pellets were removed, the marbles were wiped with alcohol, and the bedding was thoroughly mixed and smoothed before the marbles were rearranged in the same grid pattern for the next animal. Data were plotted using GraphPad Prism 9.

**Self-grooming**

The Self-grooming was conducted in a white plastic chamber (24  cm× 24 cm × 48 cm). Before each trial, the apparatus was cleaned to eliminate any residual odors. Each mouse was placed individually into the chamber and allowed to explore freely, and its grooming behavior was recorded over a 20‑min period. The total time spent grooming was manually scored using a stopwatch by an experimenter who was blinded to the genotype of the mice. Data were plotted using GraphPad Prism 9.

**Three‑Chamber Social Interaction Test**

The apparatus consisted of a white plastic box (0.75 cm thickness) with dimensions of 60 cm (length) × 40 cm (width) × 30 cm (height), divided into three chambers that were interconnected via small openings (4 cm × 4 cm) between adjacent compartments. Before each experiment, the entire apparatus and the wire cages were thoroughly cleaned to eliminate any residual odors. One day prior to testing, both the test and stimulus mice were habituated to the empty apparatus to reduce anxiety.

The test comprised two phases: the social preference test and the social novelty test. All stimulus and test mice were sex‑ and age‑matched. Stimulus mice were purchased from BIOCYTOGEN (https://biocytogen.com.cn/tool-mice) two days before each session. For the social preference test, a stimulus mouse was placed in a small wire cage located in one side chamber, while an identical empty cage was placed in the opposite side chamber. The test mouse was then introduced into the central chamber and allowed to explore freely for 10 min, during which its exploration behavior was recorded. For the social novelty test, a novel stranger mouse was placed into the previously empty cage, and the test mouse was again placed in the central chamber and allowed to explore for another 10 min. Between trials, the apparatus and cages were meticulously cleaned to remove feces and olfactory cues from the previous animal. Behavioral videos were analyzed using Noldus EthoVision XT software (https://noldus.com/ethovision), and the time spent exploring each side chamber was quantified. Data were plotted using GraphPad Prism 9.

**Barnes Maze Test**

The Barnes maze apparatus consisted of a circular white plastic board (122 cm in diameter, with a thickness sufficient to provide rigidity), elevated 80 cm above the floor and freely rotatable around its center. Twenty circular holes (5 cm in diameter) were evenly spaced around the perimeter, each fitted with an underlying groove to accommodate a removable escape box. The escape box was a small, light‑tight black plastic chamber used as a shelter from an aversive bright light. A bright light source was positioned directly above the center of the maze, and its intensity was set to maximum to serve as the aversive stimulus.

The test comprised two phases: a training phase and a probe (training) phase. One day before training, mice were habituated to the escape box and exposed to the bright light to encourage active escape‑seeking behavior. Training was then conducted for eight consecutive days, with two trials per mouse per day. At the start of each trial, the mouse was placed in the center of the maze and confined under an opaque black cylinder for 30 s to enhance light sensitivity. The cylinder was then removed, and the mouse’s behavior was recorded. If the mouse entered the escape box within 3 min, the trial was stopped, and the mouse was allowed to remain in the dark escape box for 1 min (with the overhead hole covered by a black plastic plate) as a reward. The latency to enter the escape box was recorded as the escape latency. If the mouse failed to find the box within 3 min, the trial was terminated at 3 min, and the mouse was gently guided into the escape box and rewarded with 1 min of darkness.

On day 9 (the probe phase), the escape box was removed. Each mouse was again placed in the center of the maze, confined under the opaque cylinder for 30 s, and then allowed to explore freely for 3 min. The time spent exploring the quadrant where the escape box had previously been located was measured as an index of spatial memory. All behavioral sessions were recorded and analyzed using Noldus EthoVision XT software (https://noldus.com/ethovision), and data were plotted using GraphPad Prism 9.

**Resident-intruder Test**

The intruder test was performed to evaluate social interaction ability. To further assess the social behavior of Rtel1 haploinsufficient mice, we conducted the intruder test. Briefly, an age‑ and sex‑matched unfamiliar mouse (stranger 1) was co‑housed with the experimental mouse for 24 h. The experimental mouse was then housed singly for five days to establish territoriality. One hour before testing, the experimental mouse was transferred to the testing room for habituation. The test consisted of five trials, each lasting 3 minutes with a 3-minute inter-trial interval. In the first four trials, the experimental mouse was exposed to stranger 1, and in the fifth trial, it was exposed to a novel stranger (stranger 2). The duration of sniffing behavior between the mice was recorded as a measure of social interest and novelty. The social novelty index was calculated as (T5 – T4) / (T5 + T4), where T4 and T5 represent the sniffing times in the fourth and fifth trials, respectively. All scoring was performed manually by an experimenter blinded to the genotypes of the mice. Behavioral sessions were video-recorded and analyzed using Noldus EthoVision XT software (https://noldus.com/ethovision), and the data were plotted using GraphPad Prism 9.

**Protein Purification of RTEL1**

The expression vector pcDNA3.1‑mouse‑Rtel1‑6×His was generated by inserting the mouse Rtel1 coding sequence (NM_001001882.4) into the pcDNA3.1(+) backbone. The construct was synthesized by PackGene (https://www.packgene.cn/). HEK293 cells were cultured in DMEM medium (catalog No. 11965092, Gibco) supplemented with 5 % fetal bovine serum (FBS) until reaching 90 % confluency. The cells were then transiently transfected with 2 μg of the plasmid per well in 6‑well plates using Lipofectamine™ 3000 (catalog No. L3000075, Invitrogen). After 72 h of culture, the cells were washed three times with ice‑cold 1× PBS, and the liquid was completely removed. The plates were then placed at –80 °C for 30 min to facilitate cell lysis. The cells were resuspended in lysis buffer containing 50 mM HEPES (pH 7.3, Beyotime, C0215), 250 mM NaCl, 1 mM EDTA, 0.1 % Tween‑20, and 10 % (v/v) glycerol (Beyotime, ST1348). The cell suspension was centrifuged at 12,000 × g for 10 min, and the supernatant was collected. The supernatant was then loaded onto a HisPur™ cobalt spin column (Thermo Fisher, 89969). The column resin was gently inverted several times to ensure homogeneity, and the flow‑through was allowed to pass. This step was repeated until all the lysate had passed through the column. The protein was eluted with elution buffer containing 50 mM Tris, 300 mM NaCl, and 500 mM imidazole. The eluate was concentrated using Amicon™ Ultra‑15 centrifugal filter units (UFC901008). Protein concentration was determined by UV absorbance at 280 nm. Aliquots of the purified protein were stored at –80 °C until further use.

**Western Blotting**

Proteins for Western blotting were obtained from two sources. For brain tissue samples, RIPA lysis buffer (Beyotime, P0013B) supplemented with protease inhibitors (Beyotime, P1005) was added to the tissue, and the samples were homogenized thoroughly using a tissue grinder. The homogenates were centrifuged at 12,000 × g for 10 min, and the supernatants were collected. The volume of each supernatant was recorded, and 5× loading buffer was added, followed by heating at 100 °C in a metal bath for 10 min. The resulting protein samples were separated by SDS‑PAGE using running buffer (Beyotime, P0014C) at a constant voltage of 80 V. After electrophoresis, proteins were transferred onto PVDF membranes (Thermo Fisher, 88518) using a wet transfer system at 200 mA for 2 h. The membranes were then blocked with 5 % (w/v) BSA in TBST overnight at 4 °C, followed by incubation with primary antibodies (1:1000) for 8 h and subsequently with HRP‑conjugated secondary antibodies (1:5000) for 2 h at room temperature. After three washes with 1× TBST (5 min each), protein signals were detected using chemiluminescent substrate (Thermo Fisher, 34580) on a Bio‑Rad imaging system (BIO‑RAD, 12003154).

To investigate the effects of the p.W289X and p.R734W mutations of the human RTEL1 gene on protein expression, wild‑type (WT), p.W289X, and p.R734W coding sequences (NM_001283010) were cloned into the pcDNA3.1(+) vector. The resulting constructs were transfected into HEK293 cells at 90 % confluency using Lipofectamine™ 3000. Cells were cultured in DMEM medium (95 % DMEM + 5 % FBS) for 72 h to allow sufficient protein expression. Cells were then lysed with RIPA buffer containing protease inhibitors, and the lysates were centrifuged at 12,000 × g for 10 min. The supernatants were collected, mixed with 5× loading buffer, and heated at 100 °C for 10 min. Western blotting was performed as described above. The relative protein expression levels were quantified by densitometry using ImageJ software, and data were plotted with GraphPad Prism 9.

**Immunofluorescence**

After behavioral testing, mice were transcardially perfused with ice‑cold 1× PBS to thoroughly replace the blood, followed by perfusion with 4 % paraformaldehyde (PFA) in PBS. Successful PFA perfusion was confirmed by observing twitching and upward curling of the tail during the perfusion. The brains were then carefully removed and post‑fixed in 4 % PFA overnight at 4 °C. Subsequently, brains were sequentially dehydrated in 15 % sucrose solution for 24 h and then in 30 % sucrose solution for 48 h, until the brains sank to the bottom, indicating complete dehydration. For cryosectioning, the brains were embedded in OCT compound (SAKURA, 4583) under low‑temperature conditions, frozen at –20 °C, and then sectioned at 30 μm thickness using a Leica CM1950 cryostat at –25 °C. For immunofluorescence staining, sections containing the target nuclei were selected and blocked with TBST containing 5 % BSA and 0.3 % Triton X‑100 for 3 h at 4 °C. After three washes with 1× PBS, sections were incubated with primary antibodies diluted in 5 % BSA solution overnight at 4 °C. Following three additional washes, the sections were incubated with appropriate secondary antibodies for 2 h at room temperature. After final washes, the sections were mounted and imaged using an Olympus FV3000 confocal microscope.

**Stereotaxic Injection of AAV Viruses into the PVN**

To examine whether restoring RTEL1 expression specifically in the paraventricular nucleus (PVN) rescues the autistic‑like phenotypes in Rtel1 haploinsufficient mice, we bilaterally injected AAV9‑hSyn‑DIO‑mRtel1‑3HA or control AAV9‑hSyn‑DIO‑GFP (synthesized by PackGene, https://www.packgene.cn/) into the PVN at a volume of 25 nL per side (titer: 1 × 10¹³ GC/mL). The stereotaxic coordinates were set relative to bregma (defined as the origin) as follows: ML ±0.3 mm, AP –1.0 mm, DV –4.8 mm. Viral expression was allowed for 4 weeks before behavioral testing commenced. After the behavioral battery, animals were either transcardially perfused or blood was collected for serum separation, and plasma oxytocin (OXT) levels were measured by ELISA as needed.

**Fiber Photometry Recording of OXT Signals in the PVN**

Two‑month‑old mice were anesthetized with an intraperitoneal injection of sodium pentobarbital and placed in a stereotaxic frame. AAV9/2‑hSyn‑OT was injected into the PVN using the same coordinates (ML ±0.3 mm, AP –1.0 mm, DV –4.8 mm). After the injection, a 1.25‑mm diameter optical fiber (Inper, 200 μm/NA 0.37) was implanted into the PVN. Four weeks later, OXT signals in the PVN were recorded using a fiber photometry system (Inper) with a 470‑nm laser at 50 μW output power. Recordings were performed on six mice per genotype.

**Quantitative PCR Analysis of Chromatin Immunoprecipitation (ChIP) Assays**

To quantitatively assess the binding of RTEL1 protein to the G‑quadruplex (G4) sequences within the promoter region of the OXT gene, as well as the recognition of G4 structures by the BG4 antibody, we performed quantitative PCR (qPCR) on DNA fragments obtained from immunoprecipitation (IP) experiments using anti‑RTEL1 and anti‑BG4 antibodies. The results were normalized to those obtained with control IgG antibody IP as a reference. Each IP sample was analyzed in triplicate using SYBR Green PCR Master Mix (TOYOBO, QPK‑201). Real‑time PCR was carried out on a StepOnePlus™ Real‑Time PCR System (Applied Biosystems). The relative abundance of G4 sequences in the anti‑RTEL1 and anti‑BG4 IPs over that in the IgG IP was calculated using the fold‑enrichment method and then normalized accordingly. The qPCR primers used were: forward, 5′‑TGACCAGTCATGCTGTCACC‑3′; reverse, 5′‑AGAGCCAGTAAGCCAAGCAG‑3′. All primers were synthesized by Tsingke (<https://www.tsingke.com.cn/>).

**DNA Agarose Gel Electrophoresis**

To determine whether the relevant antibodies effectively enriched target DNA fragments containing the G‑quadruplex (G4) sequences within the OXT promoter region during immunoprecipitation (IP), the DNA fragments obtained from the IP experiments were first subjected to PCR using the G4‑specific primers (forward: 5′‑TGACCAGTCATGCTGTCACC‑3′; reverse: 5′‑AGAGCCAGTAAGCCAAGCAG‑3′), Spcas9 primers (forward: 5′‑TATGGGTCGTCACAAACCGG‑3′; reverse: 5′‑TTTGCAGCTGGGTGTTTTCG‑3′). The PCR products were then analyzed by gel electrophoresis. Specifically, an appropriate volume of InstantView™ Red Fluorescent DNA Loading Buffer (Beyotime, D0076) was added to the PCR products, and the mixture was thoroughly mixed before loading onto a BeyoGel™ TBE polyacrylamide precast gel (Beyotime, D0186S), which is capable of resolving nucleic acid fragments of approximately 10 bp. Electrophoresis was performed in TBE running buffer (Beyotime, R0223) at a constant voltage of 150 V for approximately 50 min, until the bromophenol blue dye front had migrated to the bottom of the gel. The gel was then visualized using an appropriate imaging system. Because fluorescent dyes were incorporated, all steps were carried out under protection from light to prevent signal quenching.

A similar procedure was employed to distinguish G‑quadruplex (G4) secondary structures from linear DNA conformations in vitro. Briefly, the G4‑forming oligonucleotide, the G4‑mutant sequence (G4mut), and their corresponding complementary trap oligonucleotides were chemically synthesized. The synthetic G4 oligonucleotides were diluted to 25 nM in Buffer H (25 mM HEPES, pH 7.3, 100 mM KCl, 10 mM MgCl₂, 2.5 % (v/v) glycerol, and 5 mM TCEP) and then subjected to an annealing procedure to promote G4 secondary structure formation. The annealing conditions were as follows: heating at 95 °C for 10 min, followed by gradual cooling to 25 °C over a period of 3 h. The annealed products were then analyzed by DNA agarose gel electrophoresis to verify the formation of G4 structures.

**Protein–DNA Complex Electrophoresis**

To investigate the formation of protein–DNA complexes between RTEL1 and G‑quadruplex (G4) structures in vitro, the RTEL1 protein was incubated with 25 nM of the 3′‑Cy3‑labeled G4‑forming oligonucleotide (pre‑folded into G4 structures) at 25 °C for an appropriate duration to allow complex assembly. The samples were then subjected to electrophoresis on a native DNA agarose gel in TBE running buffer (Beyotime, R0223) at 150 V for approximately 50 min, until the bromophenol blue dye front had migrated to the bottom of the gel. Gels were then visualized using an appropriate imaging system. Because fluorescent dyes were incorporated, all steps were carried out under protection from light.

To examine the ATP‑dependent unwinding activity of RTEL1 on G4 structures in vitro, the RTEL1 protein was first incubated with 25 nM of the 3′‑Cy3‑labeled G4 structure at 25 °C to form protein–DNA complexes. Subsequently, 10 mM ATP and 200 nM of the complementary trap oligonucleotide (which binds to the unwound G4 strand) were added to facilitate the formation of G4‑trap duplexes. After incubation, the resulting mixture containing protein–DNA complexes, residual G4 structures, and G4‑trap duplexes was analyzed by DNA agarose gel electrophoresis.

**Chromatin Immunoprecipitation (ChIP) Assay**

To investigate whether RTEL1 protein binds to the G‑quadruplex (G4) sequences in the promoter region of the OXT gene and whether these G4 structures are formed in vivo and recognized by the BG4 antibody, we performed ChIP assays using the High‑sensitivity ChIP Kit (ab185913, Abcam). For each experiment, approximately 50 mg of brain tissue (equivalent to ~1 × 10⁶ cells) was used. Tissue samples could be snap‑frozen at –80 °C for later use. Prior to processing, the tissue was minced into 1–2 mm³ pieces and then crosslinked with 1 % formaldehyde (37 % stock diluted accordingly) at a ratio of 1 mL per 50 mg tissue. Crosslinking was terminated by the addition of glycine. The fixed tissue was then homogenized in 0.5 mL of lysis buffer per 50 mg tissue, and after resuspension, the supernatant was collected and mixed with ChIP buffer for chromatin shearing by sonication. The ChIP reaction mixture was then prepared according to the kit protocol and subjected to immunoprecipitation with the relevant antibodies. After the ChIP reaction, the DNA was purified and subsequently analyzed by PCR (or qPCR) using appropriate primers. The primary antibodies used in this experiment are BG4(MABE917, Merck), Rtel1(25337-1-AP, Proteintech), IgG(ab172730, Abcam), RANPⅡ(ab238146,Abcam)

**Circular Dichroism**

For circular dichroism (CD) spectroscopy, 10 μM G4 oligonucleotide was diluted in a buffer containing 150 mM KCl and 30 mM Tris‑HCl. The solution was then subjected to an annealing program on a PCR instrument (RWD, M2‑96SG): heated to 95 °C for 10 min, followed by gradual cooling to 25 °C over a period of 3 h. CD spectra were subsequently recorded using a 1‑cm path‑length cuvette over a wavelength range of 210–330 nm.

**Enzyme‑Linked Immunosorbent Assay (ELISA)**

To investigate the effect of Rtel1 haploinsufficiency on oxytocin (OXT) levels in the paraventricular nucleus (PVN) of mice, and to assess whether PVN‑specific restoration of Rtel1 expression could rescue OXT levels, we measured plasma OXT concentrations using an Oxytocin ELISA Kit (Sangon Biotech, D751010). After behavioral testing, blood was collected from the retro‑orbital sinus of mice into tubes containing anticoagulant. The samples were immediately centrifuged at 1,000 × g for 15 min to obtain serum, which could be temporarily stored at –80 °C for up to one month before use. OXT levels in the serum were determined following the manufacturer’s protocol provided with the kit. A standard curve was constructed by plotting the optical density (OD) values against the concentrations of the standards, and the data were fitted using a four‑parameter logistic model.

**Supplemental Figure legends**

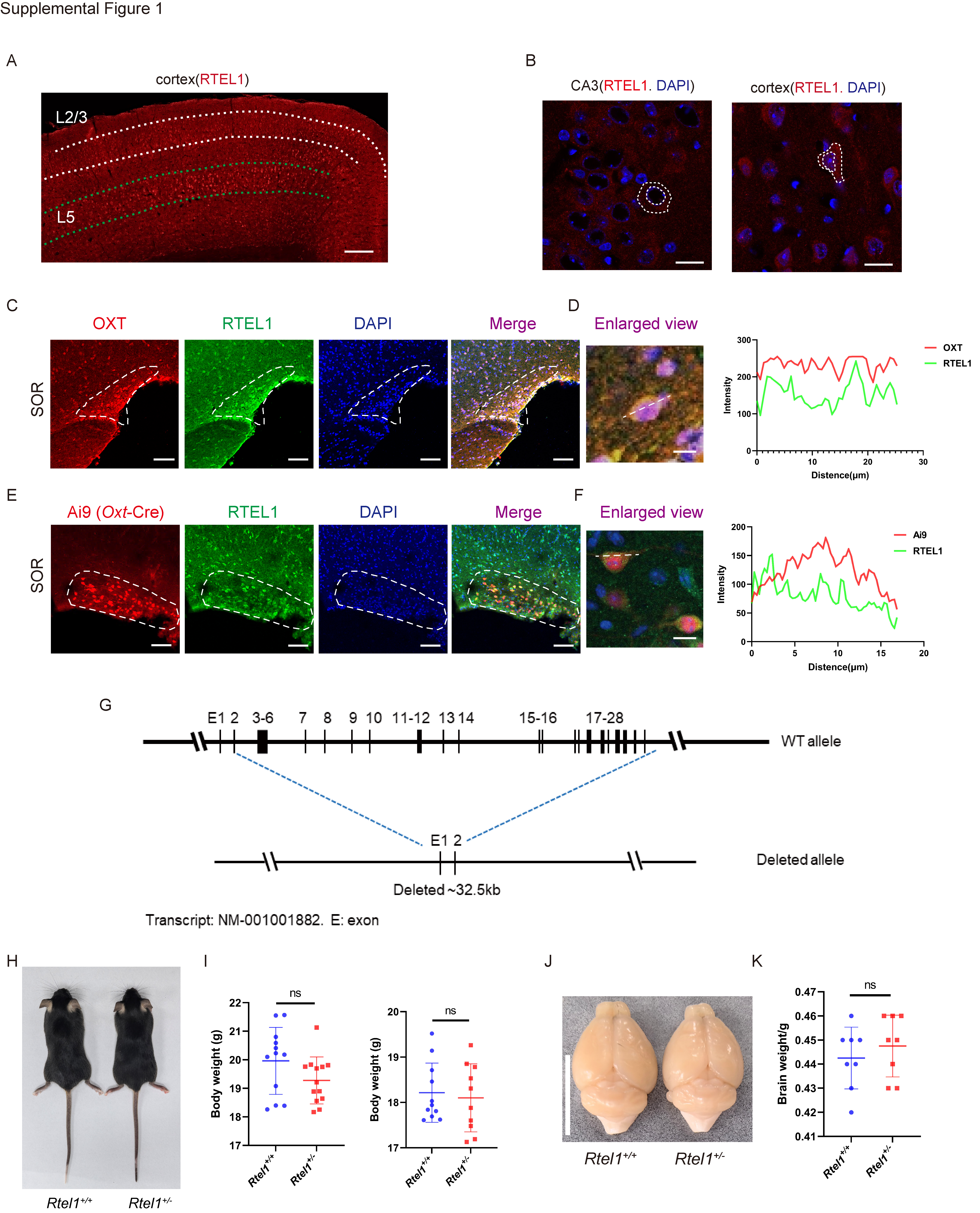

**Figure S1. RTEL1 distribution in the adult mouse brain and gross characterization of *Rtel1*^+/−^ mice. Related to Figure 1**

(A) Representative immunofluorescence image of RTEL1 in the cortex of an adult wild-type mouse. Dotted lines delineate cortical layers 2/3 and 5. Scale bar, 200 μm.

(B) Representative high-magnification images of RTEL1 (red) and DAPI (blue) in hippocampal CA3 and cortex. Dashed outlines indicate representative cells or nuclei. Scale bars, 10 μm.

(C) Representative images of OXT (red), RTEL1 (green), and DAPI (blue) in the supraoptic region, together with merged and enlarged views. Scale bars, 100 μm; enlarged view, Scale bar, 10 μm.

(D) Line-scan fluorescence-intensity profiles of OXT and RTEL1 along the line indicated in the enlarged image in (C).

(E) Representative images of the Ai9 tdTomato Oxt-lineage reporter (red), RTEL1 (green), and DAPI (blue) in the supraoptic region of Oxt-Cre;Ai9 mice, together with merged and enlarged views. Scale bars, 100 μm; enlarged view, Scale bar, 10 μm.

(F) Line-scan fluorescence-intensity profiles of Ai9 tdTomato and RTEL1 along the line indicated in the enlarged image in (E).

(G) Schematic of the wild-type *Rtel1* allele and the engineered allele containing an approximately 32.5-kb deletion spanning exons 3–28. Exon organization is shown for transcript NM_001001882.

(H) Representative dorsal images of 8-week-old *Rtel1*^+/+^ and *Rtel1*^+/−^ mice.

(I) Body weights of 8-week-old male mice (left; n = 12 mice per genotype) and female mice (right; n = 10 mice per genotype).

(J) Representative dorsal images of brains from *Rtel1*^+/+^ and *Rtel1*^+/−^ mice. Scale bar, 1 cm.

(K) Brain weights of 8-week-old male *Rtel1*^+/+^ and *Rtel1*^+/−^ mice (n = 8 mice per genotype).

Data are presented as mean ± SD. Comparisons in (I) and (K) were performed using two-sided unpaired Student’s t-tests. **p < 0.01; ns, not significant.

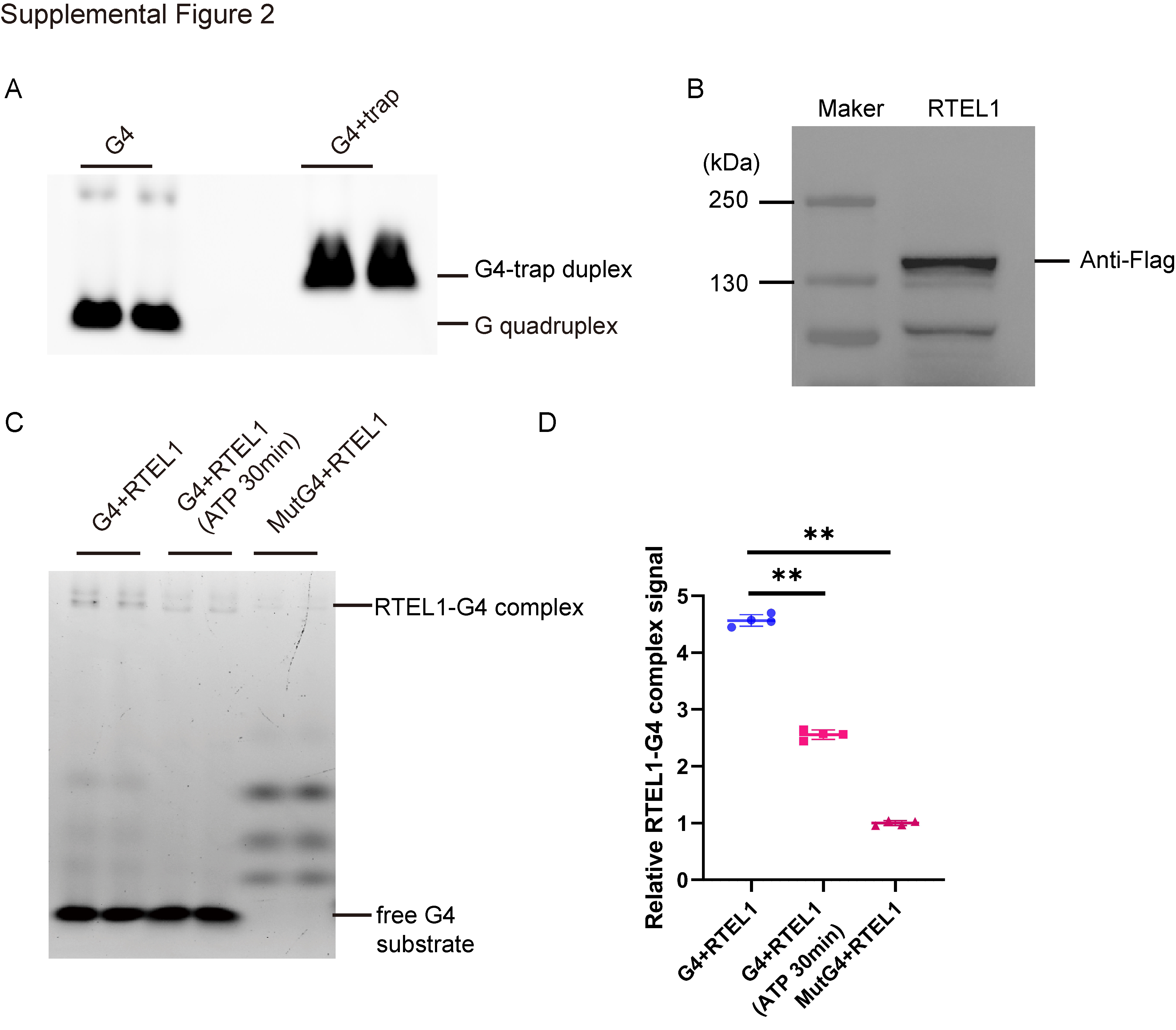

**Figure S2. In vitro formation of the Oxt promoter G-quadruplex and its interaction with RTEL1. Related to Figure 2**

(A) Representative native gel electrophoresis of the *Oxt* promoter-derived G4 oligonucleotide folded in K^+^-containing buffer, either alone or after annealing with its complementary trap strand. The positions of the G-quadruplex and linear duplex DNA species are indicated.

(B) Anti-FLAG immunoblot of the purified FLAG-tagged RTEL1 preparation used for the in vitro assays. Molecular-mass markers are indicated in kilodaltons.

(C) Representative electrophoretic mobility-shift assay of the promoter G4 oligonucleotide incubated with purified RTEL1, either in the absence of ATP or following a 30-min incubation with ATP, and of the mutant G4 oligonucleotide incubated with RTEL1. The positions of the RTEL1–G4 complex and free G-quadruplex species are indicated.

(D) Densitometric quantification of the RTEL1-associated DNA signal in (C) (n = 4 replicates).

Data are presented as mean ± SD. **p < 0.01.

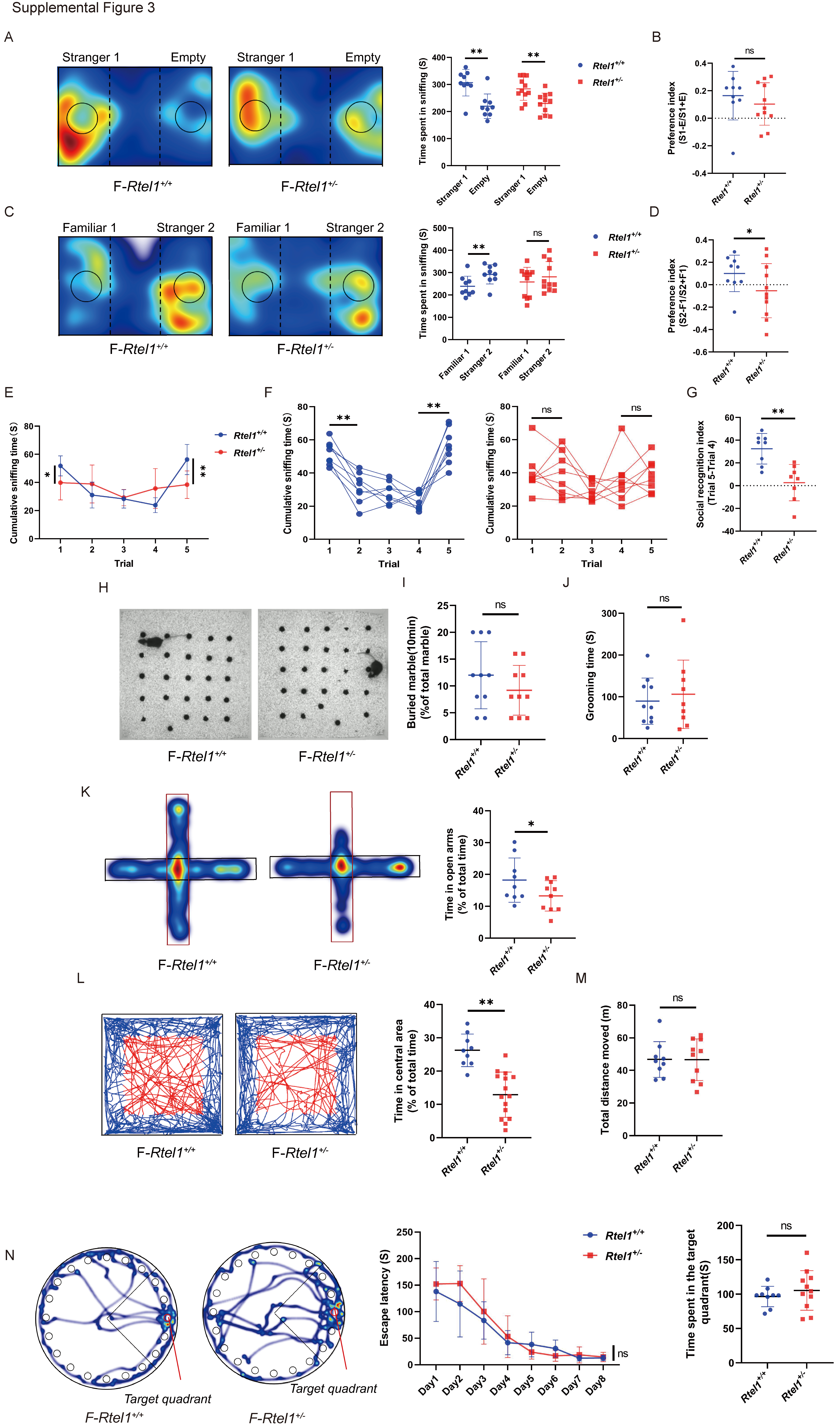

**Figure S3. Female *Rtel1*^+/−^ mice exhibit impaired social novelty and increased anxiety-related behavior. Related to Figure 4**

(A) Representative occupancy heatmaps from the three-chamber social-preference test and quantification of time spent sniffing Stranger 1 versus an empty enclosure by female *Rtel1*^+/+^ and *Rtel1*^+/−^ mice.

(B) Social-preference index, calculated as (Stranger 1 sniffing time − empty-enclosure sniffing time)/(Stranger 1 sniffing time + empty-enclosure sniffing time).

(C) Representative occupancy heatmaps from the social-novelty phase of the three-chamber test and quantification of time spent sniffing the familiar mouse (Familiar 1) versus a novel mouse (Stranger 2).

(D) Social-novelty preference index, calculated as (Stranger 2 sniffing time − Familiar 1 sniffing time)/(Stranger 2 sniffing time + Familiar 1 sniffing time). For (A)-(D), n=9 *Rtel1*^+/+^ female mice and 11 *Rtel1*^+/−^ mice.

(E) Mean sniffing time across the five-trial resident–intruder assay. Mice were repeatedly exposed to the same intruder during trials 1–4 and to a novel intruder during trial 5.

(F) Sniffing times of individual female *Rtel1*^+/+^ and *Rtel1*^+/−^ mice across the five resident–intruder trials.

(G) Social-recognition index, calculated as the difference in sniffing time between trials 5 and 4. For (E)-(G), n = 8 mice per genotype.

(H) Representative images from the marble-burying test.

(I) Percentage of marbles buried during the 10-min test. n = 10 mice per genotype.

(J) Total self-grooming duration. n=10 *Rtel1*^+/+^ female mice and 9 *Rtel1*^+/−^ mice.

(K) Representative occupancy heatmaps from the elevated plus maze and quantification of the percentage of total time spent in the open arms. n=9 *Rtel1*^+/+^ female mice and 10 *Rtel1*^+/−^ mice.

(L) Representative open-field trajectories and quantification of the percentage of total time spent in the center zone. n=9 *Rtel1*^+/+^ female mice and 15 *Rtel1*^+/−^ mice.

(M) Total distance traveled in the open-field test. n=9 *Rtel1*^+/+^ female mice and 10 *Rtel1*^+/−^ mice.

(N) Representative trajectories from the Barnes maze probe trial, escape latency across eight training days, and time spent in the target quadrant during the probe trial.

Data are presented as mean ± SD, and each point represents one mouse. *p < 0.05; **p < 0.01; ns, not significant. n=9 *Rtel1*^+/+^ female mice and 11 *Rtel1*^+/−^ mice.

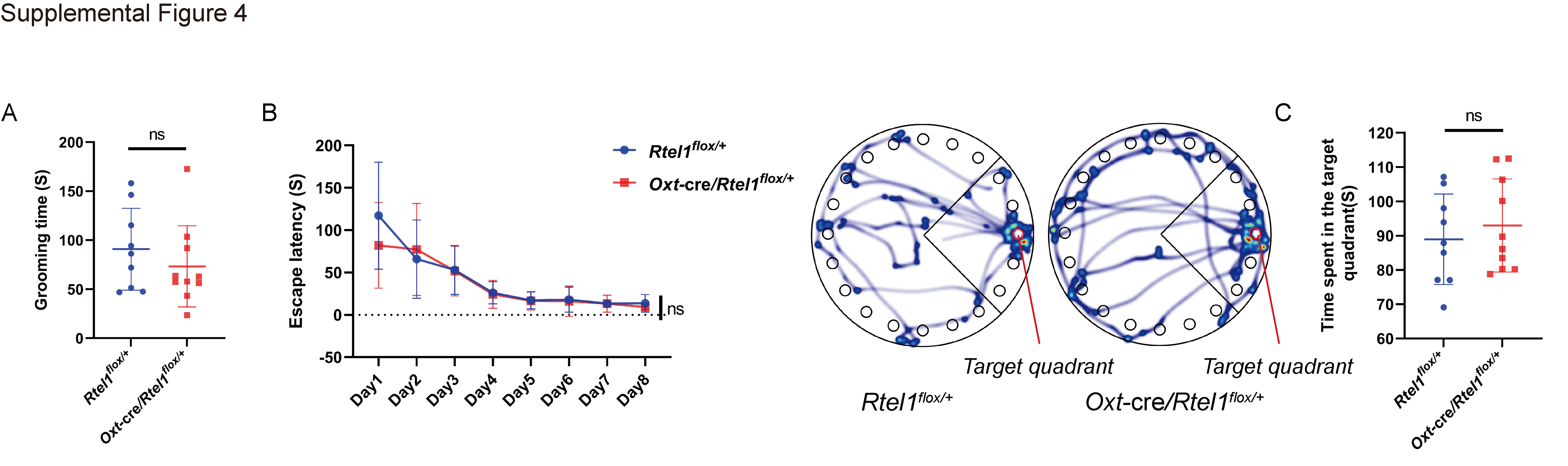

**Figure S4. Self-grooming and Barnes maze performance in male Oxt-lineage Rtel1 conditional heterozygous mice. Related to Figure 5**

(A) Total self-grooming duration in male *Rtel1*^flox/+^ control and Oxt-Cre; *Rtel1*^flox/+^ conditional heterozygous mice. n = 9 *Rtel1*^flox/+^ control mice and 10 *Oxt*-Cre; *Rtel1*^flox/+^.

(B) Escape latency across eight training days of the Barnes maze. Escape latency progressively decreased in both groups, with no significant difference between genotypes.

(C) Representative movement trajectories during the Barnes maze probe trial and quantification of the time spent in the target quadrant. For (B)-(C), n = 9 *Rtel1*^flox/+^ control mice and 10 *Oxt*-Cre; *Rtel1*^flox/+^.

Data are presented as mean ± SD, and each point represents one mouse. n = 8 male *Rtel1*^flox/+^ and Oxt-Cre; *Rtel1*^flox/+^ mice. Comparisons in (A) and (C) were performed using two-sided unpaired Student’s t tests. Training data in (B) were analyzed using a two-way repeated-measures ANOVA, with genotype as the between-subject factor and training day as the within-subject factor. ns, not significant.

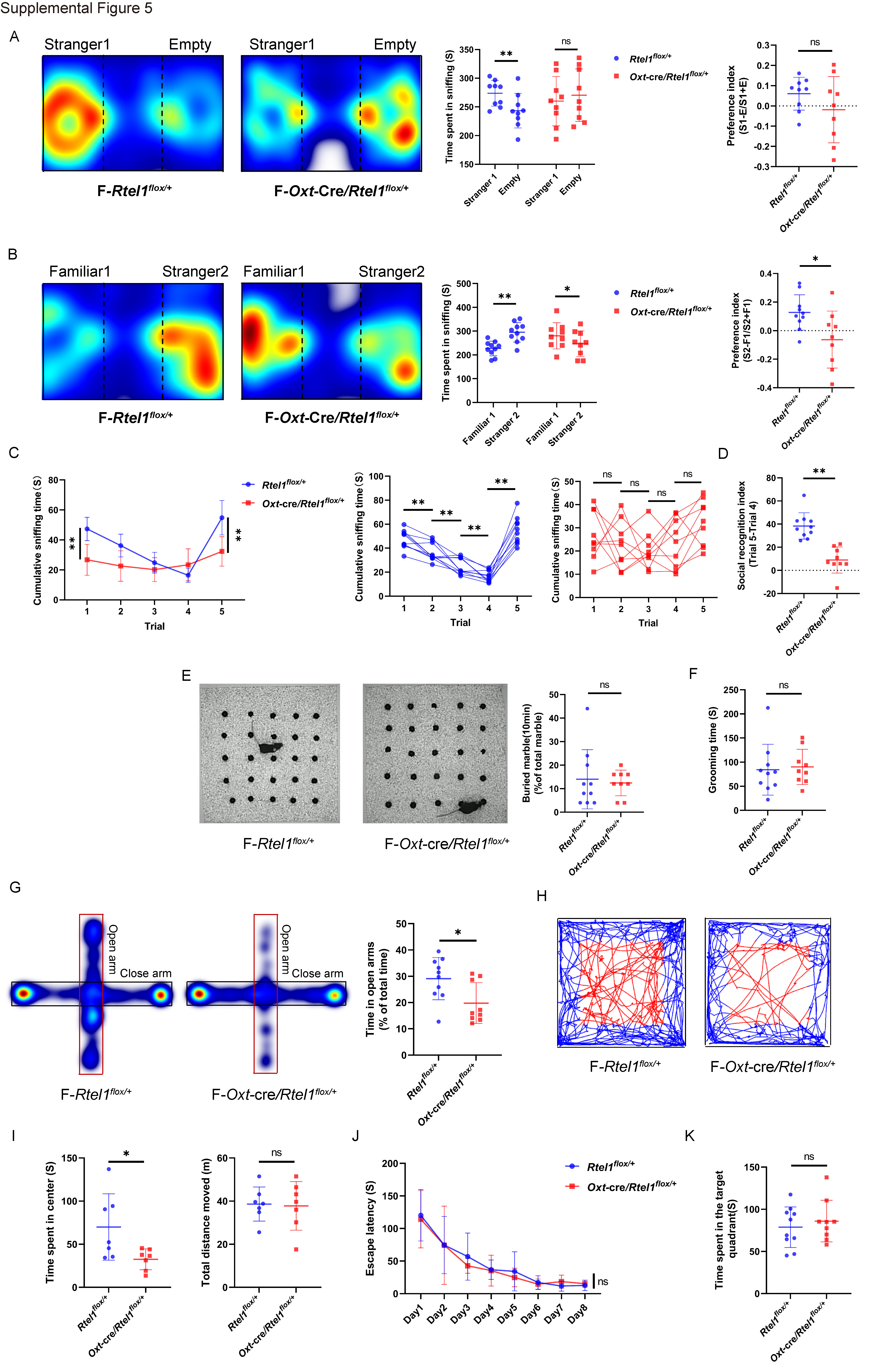

**Figure S5. Female Oxt-lineage Rtel1 conditional heterozygous mice exhibit impaired social novelty and increased anxiety-related behavior. Related to Figure 5**

(A) Representative occupancy heatmaps from the three-chamber social-preference test, quantification of time spent sniffing Stranger 1 versus an empty enclosure, and the social-preference index in female *Rtel1*^flox/+^ control and Oxt-Cre; *Rtel1*^flox/+^ conditional heterozygous mice. The index was calculated as (Stranger 1 sniffing time − empty-enclosure sniffing time)/(Stranger 1 sniffing time + empty-enclosure sniffing time). n = 9 mice per genotype.

(B) Representative occupancy heatmaps from the social-novelty phase, quantification of time spent sniffing the familiar mouse (Familiar 1) versus a novel mouse (Stranger 2), and the social-novelty preference index. The index was calculated as (Stranger 2 sniffing time − Familiar 1 sniffing time)/(Stranger 2 sniffing time + Familiar 1 sniffing time). n = 9 mice per genotype.

(C) Mean sniffing time and individual-animal trajectories across the five-trial resident–intruder assay. Mice were repeatedly exposed to the same intruder during trials 1–4 and to a novel intruder during trial 5.

(D) Social-recognition difference score, calculated as the sniffing time in trial 5 minus that in trial 4. n = 10 *Rtel1^flox/+^* control mice and 9 *Oxt*-Cre; *Rtel1^flox/+^*.

(E) Representative images from the marble-burying test and quantification of the percentage of marbles buried during the 10-min test. n = 10 *Rtel1^flox/+^* control mice and 9 *Oxt*-Cre; *Rtel1^flox/+^*.

(F) Total self-grooming duration. n = 10 *Rtel1^flox/+^* control mice and 9 *Oxt*-Cre; *Rtel1^flox/+^*.

(G) Representative occupancy heatmaps from the elevated plus maze and quantification of the percentage of total time spent in the open arms. n = 10 *Rtel1^flox/+^* control mice and 9 *Oxt*-Cre; *Rtel1^flox/+^*.

(H) Representative movement trajectories from the open-field test.

(I) Time spent in the center zone and total distance traveled during the open-field test. n = 7 mice per genotype.

(J) Escape latency across eight training days of the Barnes maze.

(K) Time spent in the target quadrant during the Barnes maze probe trial. n = 10 *Rtel1^flox/+^* control mice and 9 *Oxt*-Cre; *Rtel1^flox/+^*.

Data are presented as mean ± SD, and each point represents one mouse. *p < 0.05; **p < 0.01; ns, not significant.

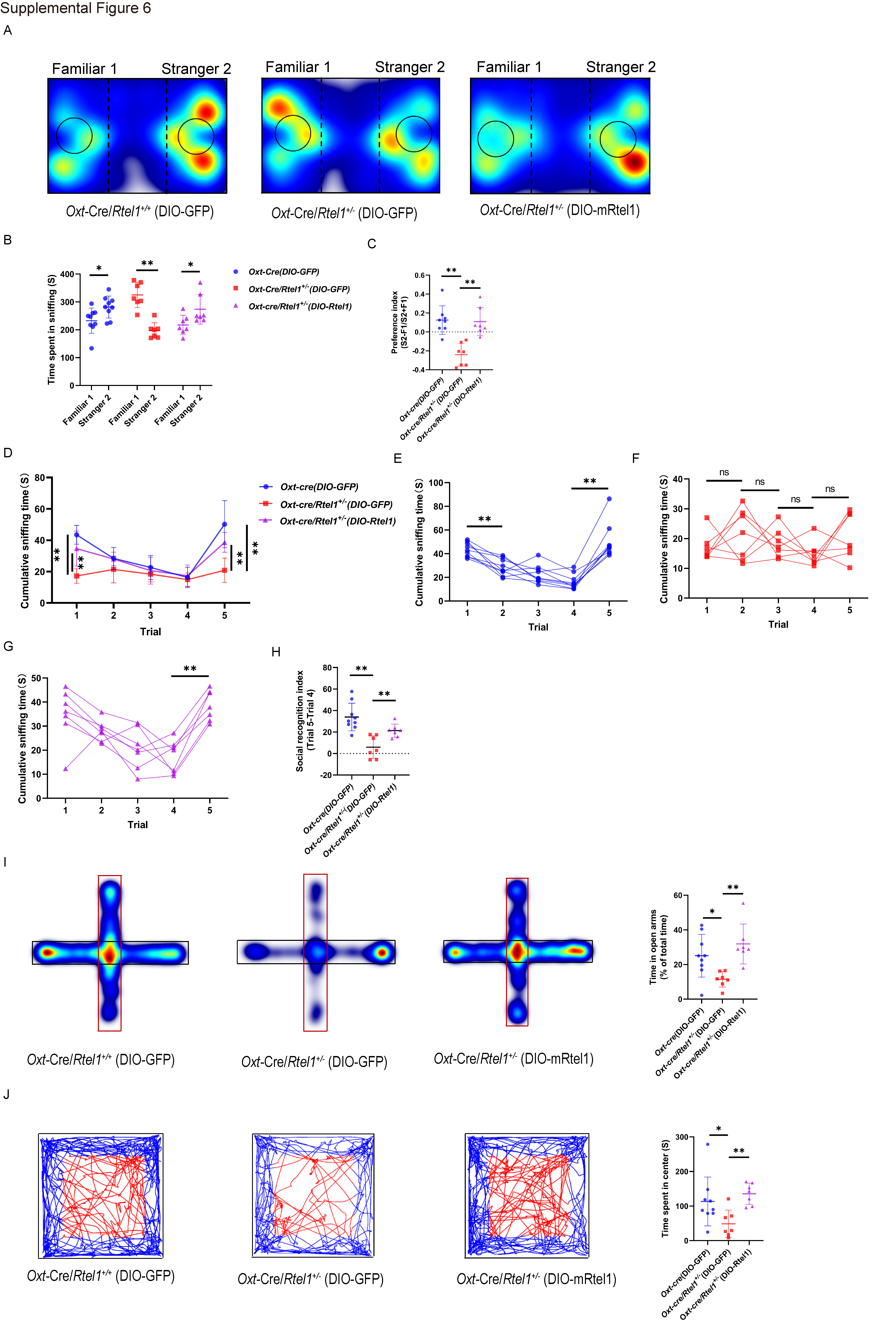

**Figure S6. PVN-targeted RTEL1 re-expression ameliorates social-recognition and anxiety-related phenotypes in female Rtel1 haploinsufficient mice. Related to Figure 6**

Two-month-old female mice received bilateral PVN injections of AAV9-hSyn-DIO-GFP or AAV9-hSyn-DIO-mRtel1-3HA and underwent behavioral testing one month later, as outlined in Figure 6A. The experimental groups were Oxt-Cre; *Rtel1*^+/+^ mice receiving DIO-GFP, Oxt-Cre; *Rtel1*^+/−^ mice receiving DIO-GFP, and Oxt-Cre; *Rtel1*^+/−^ mice receiving DIO-mRtel1.

(A) Representative occupancy heatmaps from the social-novelty phase of the three-chamber test.

(B) Time spent sniffing the familiar mouse (Familiar 1) and a novel mouse (Stranger 2) by the three experimental groups.

(C) Social-novelty preference index, calculated as (Stranger 2 sniffing time − Familiar 1 sniffing time)/(Stranger 2 sniffing time + Familiar 1 sniffing time). For (B)-(C), n = 9 WT-GFP mice, n=7 Het-GFP and 7 Het-RTEL1.

(D) Mean sniffing time across the five-trial resident–intruder assay. Mice were repeatedly exposed to the same intruder during trials 1–4 and to a novel intruder during trial 5.

(E–G) Individual-animal sniffing-time trajectories across the resident–intruder assay for Oxt-Cre; *Rtel1*^+/+^ mice receiving DIO-GFP (E), Oxt-Cre; *Rtel1*^+/−^ mice receiving DIO-GFP (F), and Oxt-Cre; *Rtel1*^+/−^ mice receiving DIO-mRtel1 (G).

(H) Social-recognition difference score, calculated as the sniffing time in trial 5 minus that in trial 4. For (D)-(H), n = 9 WT-GFP mice, n=7 Het-GFP and 7 Het-RTEL1.

(I) Representative occupancy heatmaps from the elevated plus maze and quantification of the percentage of total time spent in the open arms. n = 9 WT-GFP mice, n=7 Het-GFP and 7 Het-RTEL1.

(J) Representative open-field trajectories and quantification of time spent in the center zone. n = 9 WT-GFP mice, n=7 Het-GFP and 7 Het-RTEL1.

Data are presented as mean ± SD, and each point represents one mouse. *p < 0.05; **p < 0.01; ns, not significant.
